# A retroviral origin of vertebrate myelin

**DOI:** 10.1101/2022.01.24.477350

**Authors:** Tanay Ghosh, Rafael G. Almeida, Chao Zhao, Abdelkrim Mannioui, Elodie Martin, Alex Fleet, Ginez Gonzalez M, David H Rowitch, Katherine Stott, Ian Adams, Bernard Zalc, Nick Goldman, David A. Lyons, Robin JM Franklin

## Abstract

Myelin, the insulating sheath that surrounds neuronal axons, is produced by oligodendrocytes in the central nervous system (CNS). This evolutionary innovation, which first appears in jawed vertebrates, enabled rapid transmission of nerve impulses, more complex brains and greater morphological diversity. Here we report that RNA level expression of RNLTR12-int, a retrotransposon of retroviral origin, is essential for myelination. We show RNLTR12-int-encoded non-coding RNA binds to the transcription factor SOX10 to regulate transcription of myelin basic protein (Mbp, the major constituent of myelin) in rodents. RNLTR12-int like sequences (which we name *RetroMyelin*) are found in all jawed-vertebrates and we further demonstrate their function in regulating myelination in two different vertebrate phyla (zebrafish and frogs). Our study therefore suggests that retroviral endogenization was a key step in the emergence of vertebrate myelin.

---

Myelination, the process by which axons are invested with a myelin sheath, had a profound impact on vertebrate evolution^1–3^. By conferring the ability to transmit by rapid saltatory conduction and assisting neuronal viability by providing local metabolic support, the myelin sheath allowed axons to function over much greater lengths and hence vertebrates to attain a larger size and diversity than would have occurred in the absence of myelination. Myelination also allowed rapid conduction without needing to increase axonal diameter, enabling the packing of larger numbers of axons necessary for the evolution of complex central nervous systems (CNS). Phylogenetically, compacted myelin and genes critical to myelination such as myelin basic protein (Mbp)^2–5^ likely appeared concurrently with the emergence of jaws in vertebrates, with myelin found in the most ancient living vertebrate, the Chondrichthyes (cartilaginous fish) but not in the Agnatha (jawless fish)^1–3, 6^. Despite the significant functional advantages associated with myelination, a molecular explanation of what triggered this critical event in vertebrate evolution remains elusive.

Endogenous retrovirus (ERV) type retrotransposons are remnants of ancient retroviral sequence that persist in the genome and provide evidence of endogenization of diversified retrovirus to their vertebrate host that began several hundreds of million years ago^7^. Retrotransposons are repeated regions in the eukaryotic genome with the potential to mobilize within the genome via an RNA intermediate. The majority have lost the ability to undergo transposition due to accumulation of mutations: however, through evolution many have become ‘domesticated’ to exert gene regulatory functions by contributing to regulatory elements or being the source of functionally relevant mRNAs that can modify transcriptional networks^8–10^. Although functional retrotransposons have been implicated in the maintenance of stem cell identity^11, 12^, their role in regulating gene networks during maturation of oligodendrocytes (OLs) to form myelinating OLs is unknown.

## RNLTR12-int is a potential regulator of Myelin/Mbp

We identified retrotransposon-specific probes using the rat Affymetrix Chip and then performed meta-analysis of microarray-based gene expression data in oligodendrocyte progenitor cells (OPCs) and OLs, isolated from postnatal day 7 (P7) rat brains (Extended Data Fig. 1 a). Analysis revealed a set of protein-coding genes and retrotransposons with differential expression during OL differentiation (Extended Data Fig. 1 b, c). Differentially expressed retrotransposons included long terminal repeats (LTR), short interspersed elements (SINE) and long interspersed elements (LINE) (Extended Data Fig. 1 b-d).

To identify potential functional relationships among retrotransposon RNAs and protein-coding mRNAs in OLs we employed weighted gene co-expression network analysis (WGCNA). This revealed 13 different modules of highly co-regulated genes, which were significantly either up or down regulated in OLs (Fig. 1 a, b). The module with the highest significant association with OLs (turquoise, Fig. 1b) was overrepresented for the gene ontology (GO) term ‘myelination’ (Fig. 1c), suggesting that retrotransposon members of this module were potentially involved in regulating expression of myelination genes. Within this module, RNLTR12-int showed relatively high levels of expression in OLs as compared to OPCs (Fig. 1d), and correlated to a myelin gene co-expression network (Fig. 1e, Extended Data Fig. 1e). We undertook further experimental analysis of the relationship of RNLTR12-int with Mbp, a key component of myelin^3^.

**Fig. 1.**
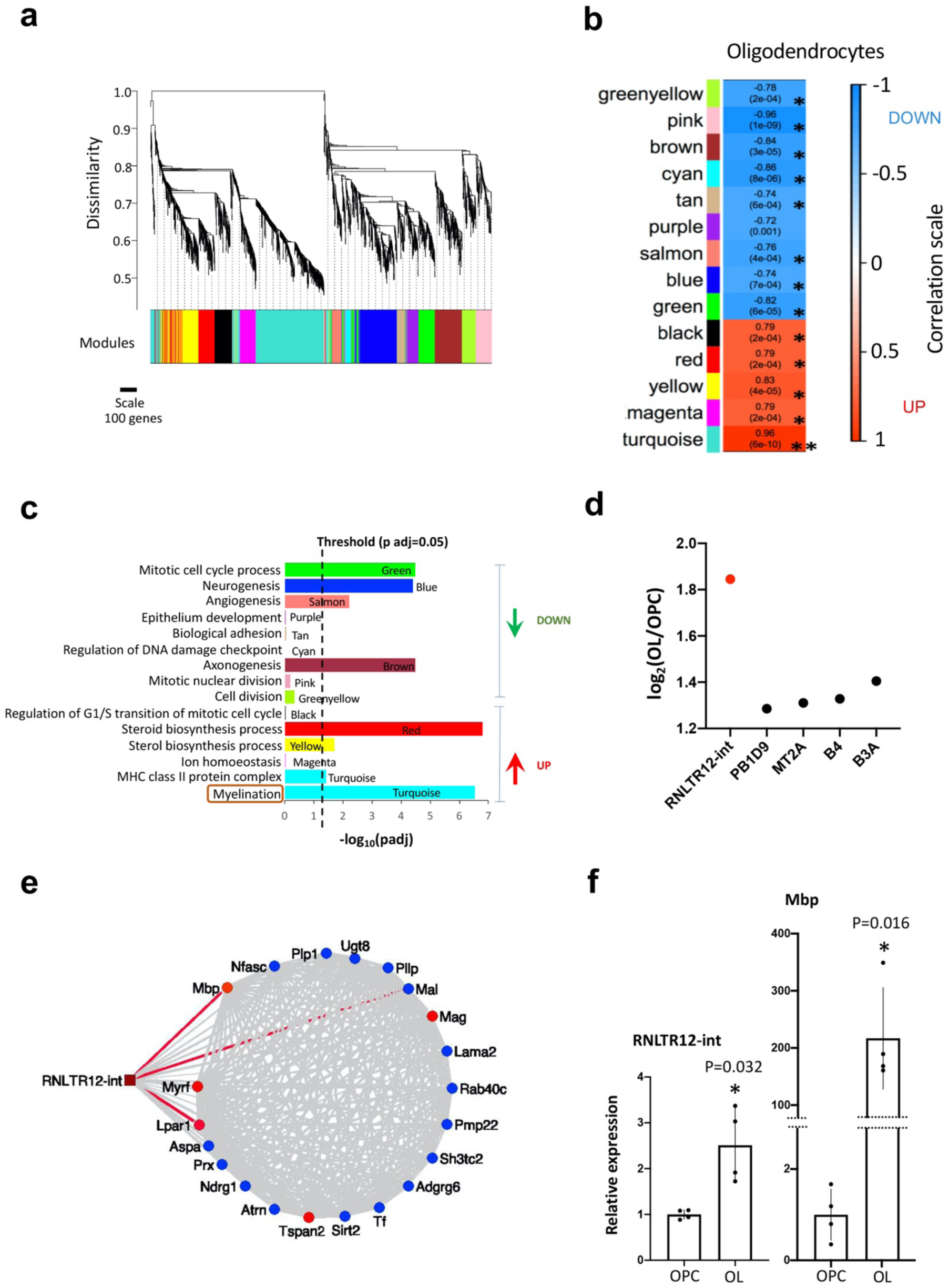
Network analyses of differentially expressed retrotransposons and protein coding genes in oligodendrocytes. **a**, Cluster dendrogram and derived modules (color labeled) after WGCNA. **b**, Association of modules with oligodendrocytes is determined by WGCNA. * p <0.0007 (Bonferroni threshold). p value in parentheses, correlation is written on top and the box is color coded according to the color scale on the left. **c,** GO-terms which are significantly overrepresented in modules. padj: adjusted p value (Benjamini-Hochberg FDR). Red and green arrow indicate the modular gene expression changes (up or down respectively) in OL as compared to OPC. **d**, Expression changes (OL vs OPC) of retrotransposons in turquoise module were plotted, as determined by microarray. P adj (Benjamini-Hochberg FDR)<0.01. **e**, Potential interaction of RNLTR12-int with myelination gene-network. Nodes: round shape, Edges: line, top 5 hubs: red round, top 3 edges: red line. Edges represent connection (expression correlation) between two nodes **f**, Expressions of RNLTR12-int transcript and Mbp in OL and OPC were determined using RT-qPCR and relative expressions as compared to OPC were plotted. Data normalized to Actb. N=4 (P7 rats), mean±SEM, *p<0.05, Student’s t test (unpaired, two tailed).

## Experimental validation of RNLTR12-int expression

To validate RNLTR12-int expression in OPCs and OLs, we extracted RNAs from Magnetic activated cell sorting (MACS)-purified OPCs (A2B5+) and OLs (MOG+) from postnatal day 7 (P7) rat brains and confirmed the expression of RNLTR12-int in OPCs and OLs by RT-PCR (Extended Data Fig. 2a). We found 2.5-fold higher levels of RNLTR12-int expression in OLs compared to OPCs by qPCR (Reverse transcription followed by qPCR, RT-qPCR) (Fig. 1f). Similarly, expression of Mbp, myelin associated glycoprotein (Mag), tetraspanin 2 (Tspan2) and PB1D9, a SINE-retrotransposon within the same WGCNA module as RNLTR12-int, were higher in OLs versus OPCs (Fig. 1f, Extended Data Fig. 2c).

## RNLTR12-int copies in the rat genome

The RNLTR12-int consensus sequence is an internal sequence of endogenous retrovirus 1 (ERV1) that has remnants of classic Gag-Pol ORFs (required for provirus replication cycle), and likely serves as a long non-coding RNA (Extended Data Fig. 2d). RNA-seq reads from OPCs were aligned to the entire consensus sequence of RNLTR12-int, confirming its full-length expression at RNA level (Extended Data Fig. 2e). In the rat genome, we identified 118 sequences of RNLTR12-int which were flanked by long terminal repeat (LTR) elements and 80 sequences which were not flanked by the LTR (Extended Data Fig. 3a, b; see methods for details). Among these 80 sequences, we found 2 genes (Pygl, Ncbp2) whose intronic regions (in opposite strand) contain fragments of RNLTR12-int. However, unlike RNLTR12-int, Pygl and Ncbp2 expression levels are not different in OPC and OL (Extended Data Fig. 3c). RNLTR12-int copies in the rat genome are less than 1% divergent from its consensus sequence. It is not clear at present whether any specific locus or multiple loci direct RNLTR12-int expression in OPC and OL.

## Mbp expression is RNLTR12-int dependent

We next examined whether the RNLTR12-int transcript (mRNLTR12-int) is required for regulation of Mbp expression in differentiating OPCs using small interfering RNA (siRNA) mediated RNA interference (RNAi)^13^ under differentiation conditions. As shown (Fig. 2a), RNLTR12-int siRNA administration inhibited expression of MBP after 5 days of differentiation (Fig. 2b) and prevented development of complex oligodendrocyte morphologies as determined by O4 immuno-staining (Extended Data Fig. 4a). We observed 98% reduction of MBP+ OLs due to inhibition of RNLTR12-int compared to controls (Fig. 2b). In contrast, the proportion (12-13%) of cells immunostained with the astrocyte marker, glial fibrillary acid protein (GFAP), were unchanged in siRNLTR12-int and control siRNA transfected samples (Extended Data Fig. 4b). RT-qPCR analyses revealed that the RNA expression of the early differentiation marker 2’,3’-Cyclic-nucleotide 3’-phosphodiesterase (Cnp) was decreased while the OPC marker platelet derived growth factor receptor alpha (Pdgfra) remained unaltered (Extended Data Fig. 4c), suggesting a role for mRNLTR12-int in differentiated cells undergoing maturation. Consistent with this, Mbp mRNA levels were drastically reduced (95%) (Fig. 2c), implying Mbp transcription is affected due to inhibition of mRNLTR12-int.

**Fig. 2.**
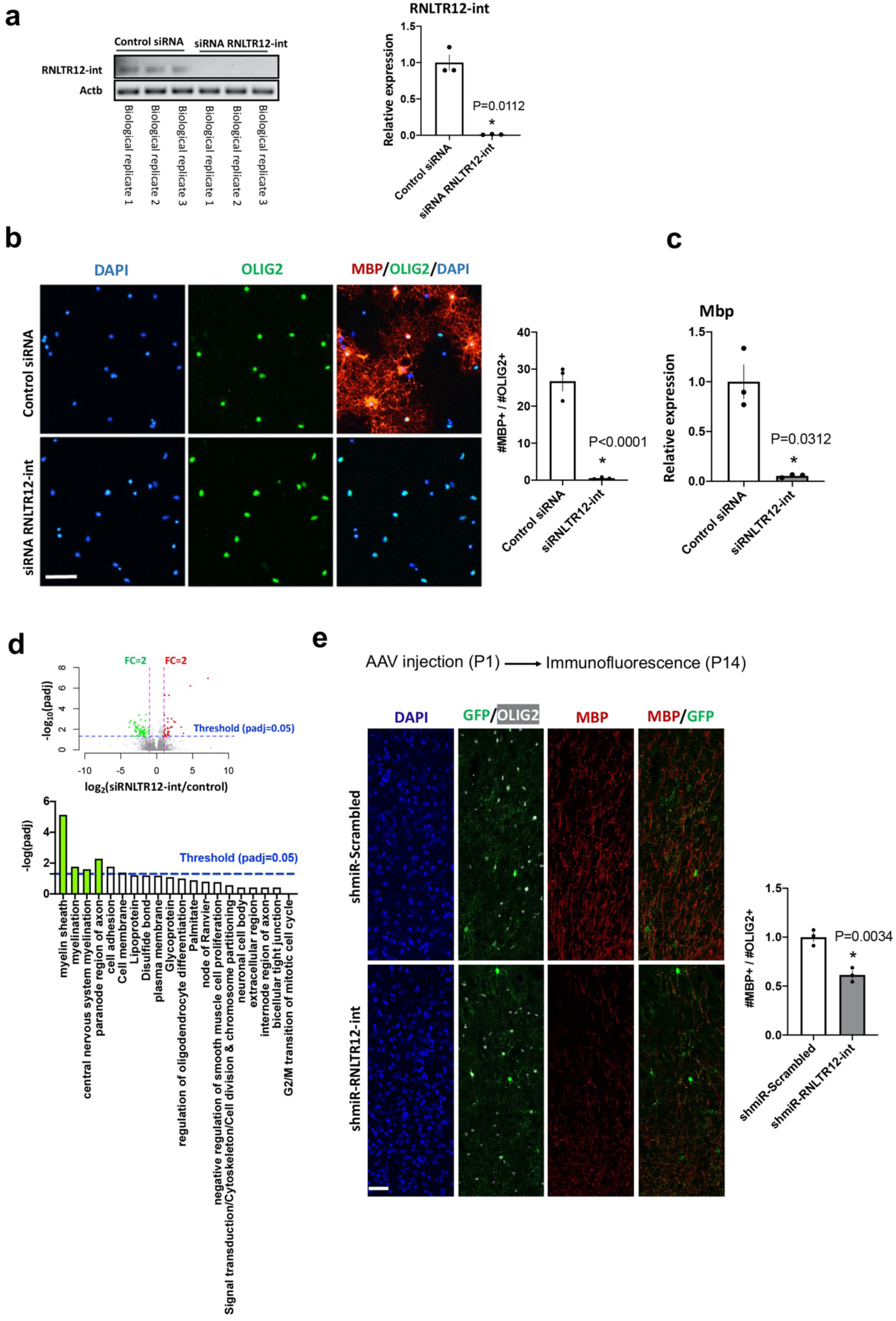
Inhibition of RNLTR12-int expression affects Mbp expression. **a**, RNLTR12-int expression was inhibited upon transfection of siRNA against RNLTR12-int (siRNLTR12-int) as determined by RT-PCR. PCR products were run in a 2% agarose gel (left). RT-qPCR analysis (right). Data were normalized to cytoplasmic b-actin (Actb). Control siRNA: siGENOME non-targeting siRNA pool (Dharmacon). N=3 independent experiments (each time 4 P7 brains were pooled for OPC isolation), mean±SEM, *p<0.05, Student’s t test (unpaired, two tailed) with Welch’s correction. **b**, Left: Immunofluorescence analysis using antibody to OLIG2 (Oligodendrocyte lineage specific marker) and to MBP (differentiation/maturation marker). Upon transfection, OPCs were maintained in differentiation media for 5 days. Scale bar: 60μm. Right: Quantification of matured OLs (MBP+, OLIG2+), presented as a percentage of total OLIG2+ cells. N=3 independent experiments (each time 3 replicates), mean±SEM, *p<0.0001, Two-way ANOVA. **c**, Effect of the inhibition of RNLTR12-int on RNA level expression of Mbp was determined by RT-qPCR. Data were normalised to Actb. N=3 independent experiments, mean±SEM, * p<0.05, Student’s t test (unpaired, two tailed) with Welch’s correction. **d**, Global changes of gene expression due to inhibition of RNLTR12-int in differentiating OPC as determined by RNAseq. Volcano plot representing differential expression (top), Up (red dot) and down (green dot) regulated genes. GO-term overrepresented in differentially expressed genes (bottom). FC: fold change, padj: adjusted p value (Benjamini-Hochberg FDR after Wald test). Genes involved in myelination (green bars) are downregulated. N=4 (control), 3 (siRNLTR12-int) independent experiments. **e**, AAV carrying SOX10-EGFP-shmiR-RNLTR12-int/Scrambled construct were injected into the deep cortex of rat at P1 (also see Extended Data Fig. 8) and brains were harvested at P14. Left: Immunofluorescence analysis using antibody to MBP, GFP, OLIG2. Scale bar: 22 μm. Right: MBP intensity were quantified and plotted relative to shmiR-Scrambled infected samples. N=3 different rats, mean±SEM, *p<0.01, Student’s t test (unpaired, two tailed).

We tested the effect of RNLTR12-int inhibition on global gene expression in differentiating OPCs by sequencing OPC samples treated with siRNA against RNLTR12-int or control siRNA. Gene ontology analyses revealed that the most differentially expressed genes related to myelination and were downregulated in the knock-down samples (Fig. 2d). Since RNLTR12-int fragments might exist inside a gene, we tested whether our results might have arisen due to off-target effects. A 125 bp region, flanked by qPCR primers to amplify RNLTR12-int including our siRNA target, was used as an input in BLAT (using rat genome) and annotations extracted for all BLAT hits. No BLAT hits fell within the exonic region of any genes. Some hits fell within intronic regions in 19 genes: however, none of these were differentially expressed by inhibition of RNLTR12-int (Extended Data Fig. 5a). Restoring expression of RNLTR12-int following RNLTR12-int siRNA silencing significantly elevated MBP expression in differentiating OPCs (Extended Data Fig. 5b). Together, these data supported our conclusion that the effects of RNLTR12-int inhibition were indicative of a direct role in myelination.

## RNLTR12-int regulates Mbp expression in developmental myelination

We next examined the effect of RNLTR12-int inhibition in the oligodendrocyte lineage during developmental myelination. We used a SOX10-driven shmiR construct where shRNA against RNLTR12-int is embedded into a microRNA cassette with a lineage-specific Emerald GFP (EGFP) reporter (Extended Data Fig. 6a). We used adeno associated virus (AAV) carrying SOX10-EGFP-shmiR (Extended Data Fig. 6a) *in vitro* to confirm it reliably infected and inhibited *RNLTR12-int* expression in differentiating OPCs (Extended Data Fig. 6b). We then injected AAV SOX10-EGFP-shmiR into the deep cortex of postnatal day (P) 1 rat (Extended Data Fig. 6a) and subsequently harvested brains at P14. We found that all EGFP immuno-positive cells (total GFP+ cells counted: 956 for shmiR-scrambled; 773 for shmir-RNLTR12-int) were also OLIG2 positive (Extended Data Fig. 6c), indicating oligodendrocyte lineage fidelity. Our protocol infected approximately 43% of OLIG2+ cells (956 GFP+OLIG+ out of 2214 OLIG+ cells counted in shmiR-scrambled; 773 GFP+OLIG2+ out of 1758 OLIG2+ cells counted for shmir-RNLTR12-int) (Extended Data Fig. 6c). We found an equal proportion (425 CC1+GFP+ out of 574 GFP+ cells in shmir-scrambled; 409 CC1+GFP+ out of 561 GFP+ cells in shmir-RNLTR12-int) of EGFP+ cells were immunoreactive for CC1 (an early differentiation marker of OLs) in both shmiR-RNLTR12-int and shmiR-Scrambled cases (Extended Data Fig. 6d), indicating that differentiation of OPCs into oligodendrocytes was not impaired. However, we observed a significant reduction of MBP expression in shmiR-RNLTR12-int infected brains (Fig. 2e) suggesting that MBP expression in vivo is RNLTR12-int dependent.

## RNLTR12-int transcript is essential for SOX10 binding to Mbp promoter

Because non-coding RNA also binds to certain transcription factors to affect target gene expression^14–19^, we tested if SOX10 mediated transcription of Mbp^20–21^ is regulated by RNLTR12-int. To examine if SOX10 bound to mRNLTR12-int *in vivo*, we used a SOX10 antibody and performed RNA immunoprecipitation (RIP) on post-natal rat brains. RT-qPCR analysis on RIP samples indicated that SOX10 bound mRNLTR12-int *in vivo* (Fig. 3a), and we confirmed direct binding between SOX10 and mRNLTR12-int using surface plasmon resonance analysis (Fig. 3b). We next asked whether the binding of SOX10 to the Mbp promoter is RNLTR12-int-dependent. As illustrated in Figure 3c, two conserved sequence elements (S2, S1)^21^ located 153 bp apart on the Mbp promoter are important for SOX10 binding. siRNLTR12-int transfected OPCs were allowed to differentiate for 4 days in culture. Subsequently chromatin immunoprecipitation (ChIP) with SOX10 antibody and qPCR analysis was performed. We identified a significant enrichment of the Mbp promoter region in control siRNA transfected samples (containing SOX10 immunoprecipitant), confirming SOX10 localization to Mbp promoter region comprising S2 and S1 (Fig. 3c). However, there was no enrichment of the Mbp promoter region in siRNLTR12-int transfected samples, suggesting mRNLTR12-int is essential for SOX10 binding to the Mbp promoter (Fig. 3c, d).

**Fig. 3.**
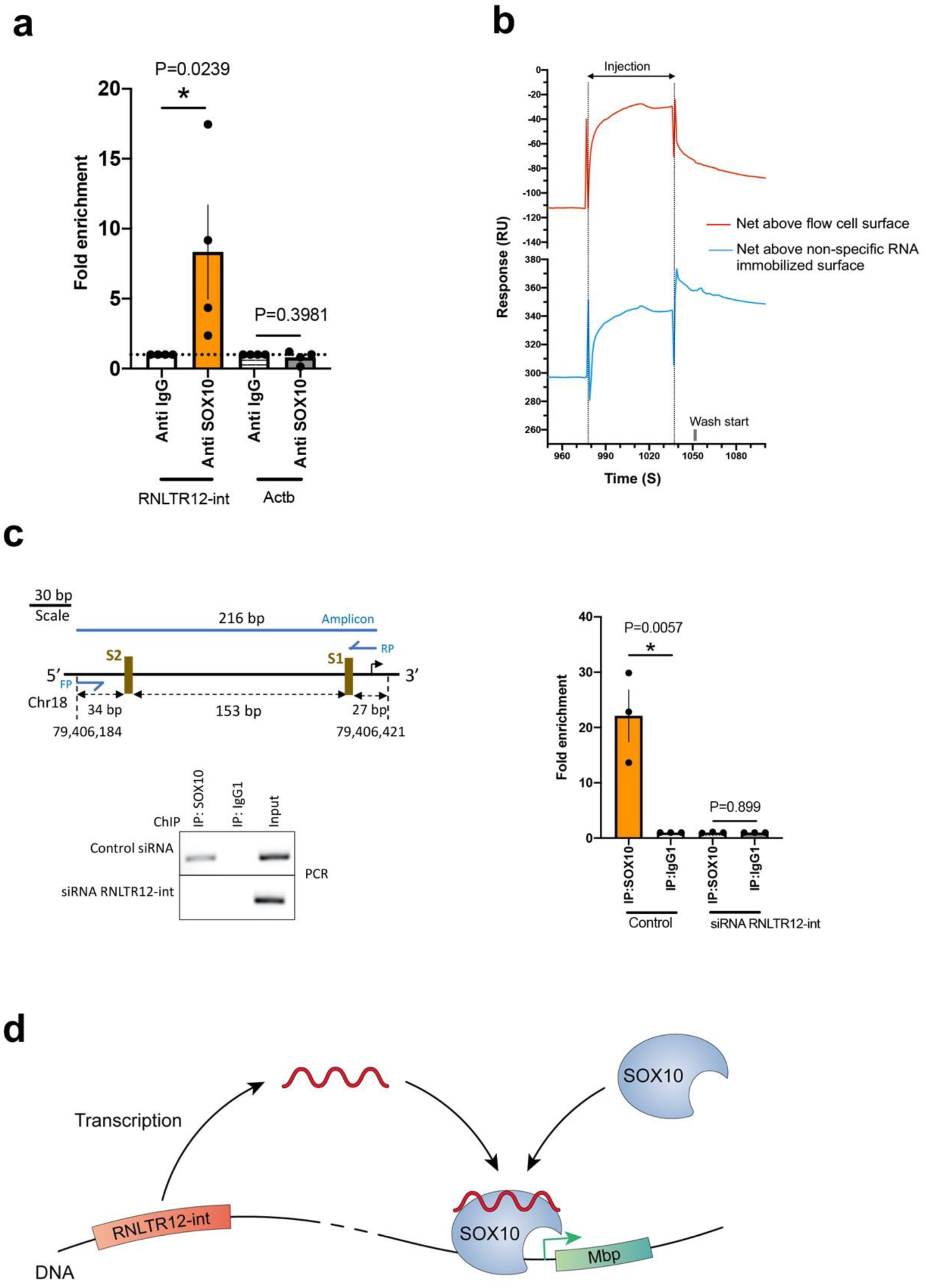
SOX10 binds to RNLTR12-int encoded transcript and this is required for SOX10 binding to Mbp promoter. **a**, RT-qPCR after crosslinking RIP (from P7 rat brains) with SOX10 or control immunoglobulin G1, showing SOX10 association with RNLTR12-int transcript. N=4 independent experiments (each time 5-6 brains were pooled). mean±SEM, * p<0.05, Ratio paired t-test (two-sided). **b**, Direct binding of RNLTR12-int transcript and SOX10, determined by surface plasmon resonance. SOX10 protein was injected over the streptavidin-coated flow cell sensor surface immobilized with 3’ end biotin-labelled RNLTR12-int/non-specific RNA or kept blank. Sensorgram represents the data corrected for non-specific binding to the flow cell surface (red) or for non-specific binding to the flow cell surface immobilized with non-specific RNA (Blue). Net positive response above these backgrounds indicates direct binding. **c**, ChIP analyses with SOX10 or control immunoglobulin G1, 4 days after siRNA transfection, showing SOX10 occupancy to Mbp promoter is affected in absence of RNLTR12-int transcript. Left: (Top) schematic representation of Mbp promoter. Rat genome assembly: rn6, FP: forward primer, RP: reverse primer for PCR amplification after ChIP. SOX10 binding conserved elements: S2 (‘AACAAT’) and S1 (‘TTCAAA’). Black right arrow after S1: Transcription start site (TSS). (Bottom) PCR amplified immunoprecipitated samples, run on 2% agarose gel. Right: qPCR analysis after ChIP. N=3 independent experiments. mean±SEM, * p<0.01, Ratio paired t-test (two-sided). **d**, Illustration represents SOX10 mediated transcription of Mbp is RNLTR12-int transcript dependent. Copies of RNLTR12-int exists in the rat genome. For simplicity reason only one is drawn. RNLTR12-int is transcribed. Association of the transcription factor SOX10 to Mbp promoter requires a direct binding interaction between SOX10 and RNLTR12-int transcript. Green arrow: TSS.

## RNLTR12-int like sequence in jawed vertebrates

Distribution of transposons in a specie’s genome is not random^22^: insertions which confer survival fitness to their host are preferentially retained through natural selection. Compacted myelin or MBP expression is evident only in jawed vertebrates and among these the most ancient living animals are the cartilaginous fishes^1–3, 6^. After establishing a direct regulatory relationship between Mbp and mRNLTR12-int in mammalian CNS, we searched for RNLTR12-int-like sequences within jawed vertebrates, jawless vertebrates and selected invertebrates based on a profile hidden Markov model, followed by repeat annotation and repeat family identification. We were able to detect RNLTR12-int-like sequences (which we have named *RetroMyelin* (<u>Retro</u>transposon sequences associated with <u>Myelin</u> evolution) in all the jawed vertebrate phyla analysed, including the ancient cartilaginous fishes (elephant sharks, whale sharks) (Fig. 4a). In contrast, we found no such sequence in jawless vertebrates (e.g., lamprey), fish-like jawless chordates (lancelet) and invertebrates (*Drosophila, C. elegans,* Sea anemone, Sea urchin) (Fig. 4a). *RetroMyelin* in jawed vertebrates all belonged to the ERV1 family, every occurrence being annotated as an internal sequence of ERV1. Furthermore, we took *RetroMyelin* sequences from human, mouse, rat, zebrafish, and elephant shark and employed the NCBI BLASTN search engine to query expressed sequence tags (ESTs) databases. We found ESTs representing these sequences with a full coverage, suggesting that they are indeed expressed at RNA-level (Extended Data Fig. 7).

**Fig. 4.**
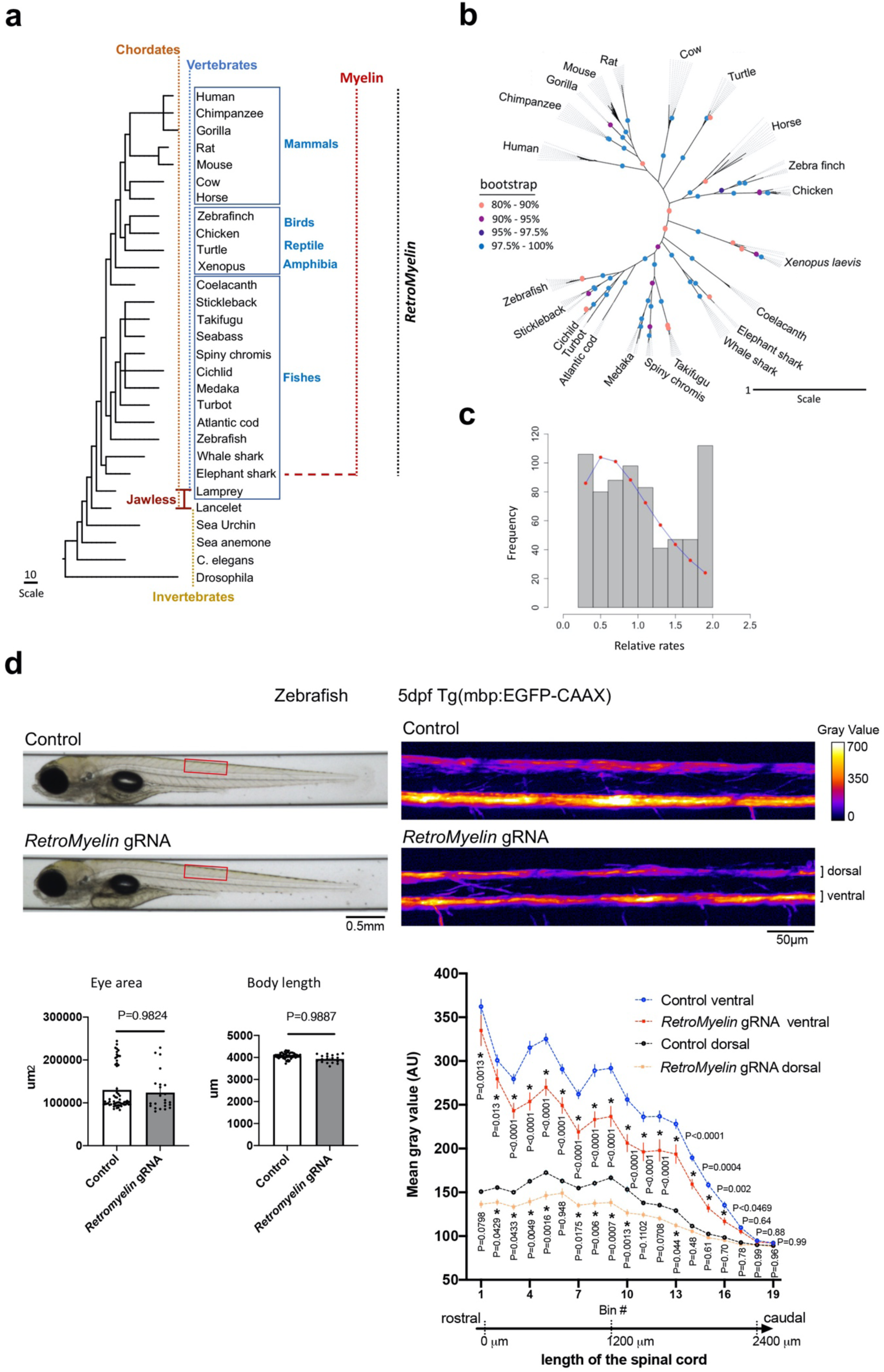
Phylogenetic relationship of RNLTR12-int and Mbp. **a**, Common Tree display represents a hierarchical view of evolutionary relationships among taxa and their lineages. Appearance of Myelin/MBP is associated with the gain of hinge-jaw in vertebrates^1–3, 6^. Presence of myelin/MBP^1–3, 6^ in most ancient living vertebrate (the cartilaginous fishes) (red dotted horizontal line) to higher vertebrates (red dotted vertical line) is marked. Black dotted vertical line marks the vertebrate phyla wherever *RetroMyelin* is identified in our study. All identified *RetroMyelin* belong to the ERV1 family, carrying internal sequence of ERV1. **b**, Phylogenetic tree (inference based on maximum likelihood and and bootstrapping) was reconstructed from the copies of *RetroMyelin* sequences from each species (detail in supplementary methods). Copies of *RetroMyelin* in each species were clustered together, with high bootstrap support values throughout (coloured circles), and particularly at host-species-defining branches. **c**, Evolutionary rate heterogeneity analysis of *RetroMyelin*. Frequency of site-wise relative rates is represented as a histogram (grey bars). Expected values of relative rate per bin were calculated from the cumulative distribution function of the inferred gamma distribution (the inferred shape parameter (α) = 2.224, standard error = 0.156) and plotted (red dots). **d**, Reduction of Mbp-promoter driven EGFP expression upon genome editing of a small region of *RetroMyelin* sequences in a transgenic zebrafish reporter line (Tg(mbp:EGFP-CAAX). left: Automated imaging of control and *RetroMyelin* gRNA injected zebrafish larva at 5 days postfertilization shown no difference in larva morphological development analysed by eye area and body length. Adjusted P value after Two way ANOVA and Benjamini-Hochberg FDR post test, mean±SEM, N=52-60 (control), 19-23 (*RetroMyelin* gRNA) animals; right: significant decrease in myelination, analysed by measuring the fluorescence intensity along the dorsal and ventral tracts of the spinal cord. * adjusted p < 0.05, Two way ANOVA (Benjamini-Hochberg FDR post test), mean±SEM, N=60 (control), 23 (*RetroMyelin* gRNA) animals.

## Acquisition of *RetroMyelin* in host genome

*RetroMyelin* sequences in vertebrate genomes could have been acquired either once before their speciation from the most recent common ancestor, or through multiple retroviral invasions after speciation had begun. In the former case, sequence divergence would have started at the ancestor stage and copies randomly sampled from a species would not be clustered together in a phylogenetic tree (Extended Data Fig. 8). However, if *RetroMyelin* were acquired after speciation then the sequence copies of each species would form separate cluster as divergence would only happen within a species. To test this, we sampled multiple *RetroMyelin* copies from 22 different species. In the reconstructed phylogenetic tree, we found that the copies from each species are clustered together, suggesting the occurrence of separate invasion events after final speciation followed by diversification of sequence copies within that host species (Fig. 4 b). Thus, the acquisition of *RetroMyelin* sequences into host specie’s genomes and their co-option for Mbp expression likley happened through convergent evolution.

Selective constraints during evolution generally limit the rapid divergence of essential genomic sequences. We employed a mutational rate heterogeneity model with the assumption that the variation of evolutionary rates of nucleotide substitution over sites followed a gamma distribution^23^. Presence of mutational rate heterogeneity was confirmed by a likelihood ratio test (2ẟ = 1528.5; p < 0.005), providing strong evidence of rate variation over sites in *RetroMyelin*. Evolutionary relative rate estimates among the sites in *RetroMyelin* suggested that there are many sites (52%) that are relatively conserved or were slowly evolving (relative rate<1) (Fig. 4c). Together, these data are suggestive of selective constraints acting on some parts of these sequences during evolution.

## RetroMyelin regulates myelination in non-mammalian classes of vertebrate

To test whether *RetroMyelin* played a similar role in the regulation of Mbp in other vertebrate classes, we employ zebrafish (*Danio rerio*) and frog (*Xenopus laevis*). We used a zebrafish reporter line Tg(mbp:EGFP-CAAX), where EGFP fluorescent intensity is a proxy measure of Mbp promoter activation. Using CRISPR/Cas9 technology, we acutely disrupted a small region of *RetroMyelin* sequences in the zebrafish genome by injecting fertilized eggs with a *RetroMyelin* guide crRNA and Cas9 enzyme (Fig. 4d, Extended Data Fig. 9). No difference of morphological development (eye area and the body length) was observed at 5 days post-fertilization (dpf) in zebrafish larva that had been injected with a *RetroMyelin* guide crRNA compared to Cas9-only control animals (Fig. 4d). However, a significant reduction of EGFP fluorescence intensity was detected along the dorsal and ventral tracts of the spinal cord (Fig. 4d), suggesting the involvement of *RetroMyelin* in regulating Mbp transcription in fish. Furthermore, *RetroMyelin* disruption in frog resulted in a significant reduction of myelin/MBP expression in tadpoles without altering oligodendrocytes number in the brain stem (Extended Data Fig. 10). Together our experiments suggest a conserved function of *RetroMyelin* in regulating Mbp transcription between fish, amphibians and mammals.

## Discussion

Generation of myelin by oligodendrocytes is not only important in development, normal physiology^24, 25^ and in regeneration following demyelination^26, 27^ but also played a central role in evolution^1–3^. We provide evidence that transcriptional regulation of Mbp is critically governed by *RetroMyelin* encoded transcripts. MBP is essential for myelination of the vertebrate CNS^2–5^, unlike the other myelin proteins MAL, MAG, CNP, PLP, PMP22, TSPAN2 that are evolutionarily older and exist in non-myelinated jawless vertebrates and invertebrates^3^. In our phylogenetic analysis, we found *RetroMyelin* exists in all jawed vertebrates where myelination is evident. We propose that in vertebrate species, ERV1 type of retrotransposons originate due to germline invasion and genomic integration of retroviruses carrying an RNLTR12-int like sequence. Separate invasion events might have happened after final speciation. The newly invaded ERV genome was either purged from the population pool (which may account for the absence of *RetroMyelin* in jawless vertebrates), or, diversification of the introduced sequence copies within each host occurred and was eventually exapted to regulate Mbp through convergent evolution. The role for ERV1 class retrotransposons in Schwann Cells, which myelinate the peripheral nervous system, remains to be addressed. Our study suggests that *RetroMyelin* was co-opted to regulate transcription of Mbp and thus endogenization of ERV1 into the vertebrate genome is coupled to the evolutionary emergence of myelination.

## Acknowledgments

We thank Peter Humphreys and Darran Clements for the technical assistance and Prof. Travis J Wheeler (University of Montana) for helpful discussion.

## Author contributions

Conceptualization: T.G., R.J.M.F.; Formal analysis: T.G., N.G.; Investigation: T.G., C.Z., R.G.A., A.M., E.M., A.F., G.G.M., K.S.; Methodology: T.G., R.G.A., R.J.M.F., D.A.L., B.Z., N.G., I.A.; Project administration: T.G., R.J.M.F.; Supervision: R.J.M.F., T.G.; Writing – original draft: T.G., R.J.M.F.; Writing – review & editing: T.G., R.J.M.F., D.A.L., N.G., D.H.R, C.Z., I.A., B.Z., A.M., R.G.A., K.S., G.G.M., E.M.

## Data and materials availability

All data are available in the main text or the supplementary materials.

## Methods

### Retrotransposable element annotation from Affymetrix microarray

Retrotransposon probes in Affymetrix Rat Genome 230 2.0 were identified and annotated as described previously^28^. Affymetrix Rat Genome 230 2.0 contains 31,000 probesets. Probe sequence were used for BLAT search in the rat genome (rn4, Nov. 2004, version 3.4) and multimapping probes were identified. Genomic co-ordinates of these probes were then compared with Repeatmasked region (rn4 release rat genome) by querying repeat masker database^29^ and annotation (class, family and element) were tabulated.

### Probe level analyses of Affymetrix microarray data

Raw data with accession number GSE11218 and GSE5940 (using Affymetrix expression GeneChip Rat Genome 230 2.0 and 230A, respectively) were extracted from Gene expression omnibus (GEO). Raw data from both the platform comprise 9 OPC and 8 OL samples. Probe level information was extracted using R library rat2302probe (for Rat genome 230 2.0) and rae230aprobe (for Rat genome 230A). Unique probe sequence in 230 2.0 and 230A platform were retained and the intersect probes (90507 in number) between two arrays were used for analysis. Background subtracted raw data were quantile normalized. Probes for which intensity were greater than sample median level in at least half the arrays for one experimental condition in a dataset were considered present^28^; absent probes were removed. Further differentially expressed probes between OL and OPC were identified by using R Bioconductor package limma^30^. P-values were adjusted to correct for multiple testing using FDR (Benjamini-Hochberg). Probes for which the difference between groups were ≥ 2 fold and the adjusted p-value < 0.05 were considered differentially expressed.

### Weighted gene co-expression network analysis (WGCNA)

WGCNA was performed by using functions^31^ available as a R package (1.20 library) installer. Because each gene is represented by multiple probes in microarray, aggregate^32^ function was used to collapse probe with median expression. Pearson correlation matrix (preserving the sign of the correlation coefficient) of pairwise comparison of gene expression levels was generated and transformed to an adjacency matrix by using soft threshold power=16 for fitting with approximate scale free topology; thereafter a topological overlap matrix (TOM) was generated. Topological overlap (TO) based dissimilarity (1-TO) measure was used for generating average linkage hierarchical clustering and cutreeDynamic function was used to identify modules. Module eigengene (ME) (i.e. 1^st^ principal component obtained after singular value decomposition) is a summary of the gene expression levels of all genes in a module. Binary numeric variables called ‘oligodendrocytes’ was generated to define OPC (=0) and OL (=1). Correlation co-efficients (Pearson) between MEs with oligodendrocytes and p-values (Student asymptotic) of corresponding correlations were obtained by using WGCNA functions. Bonferroni threshold level of significance (0.01/14 = 0.00071) was set based on the number of modules (=14).

### Overrepresentation analysis

DAVID (the Database for Annotation, Visualization and Integrated Discovery)^33^ functional annotation tool was used for analyzing GO (Gene ontology) terms overrepresented in WGCNA modules. Benjamini-Hochberg adjusted p value <0.05 was considered significant.

### Coding potential determination

Coding and non-coding potential was evaluated using CNIT (Coding-Non-Coding Identifying Tool) program^34^.

### Aligning RNA-seq reads to RNLTR12-int consensus sequence

RNAseq raw data from OPC (GSM3967160) were pre-processed through sortMeRNA^35^ to filter out reads from ribosomal RNA, adapter was trimmed, and quality paired reads were extracted using Trimmomatics^36^. RNLTR12-int consensus sequence (from Repbase-GIRI) was indexed and reads alignment to RNLTR12-int was performed by STAR_2.5.2b^37^. Subsequently sorted bam file was generated, and depth was calculated using samtools^38^.

### RNA-seq with siRNLTR12-int

OPC were isolated from P7 rats and maintained in culture media (see the section: Cultute and differentiation of OPC). Approx. 40,000 cells were plated in a 12 well plate. Subsequently transfection was performed (see the section: Transfection) and cells were kept at differentiating media for 4-5 days. 4 (control) to 3 (siRNLTR12-int) independent biological replicates were prepared (each time 4-5 P7 rat brains were pooled for OPC isolation). RNA was isolated using mirVana RNA isolation kit (Ambion). Subsequently, DNAase treatment and purification was performed and RNA quality was verified in a bioanalyzer before used for RNAseq library preparation. RIN values were: 9-10. Qiaseq Fast Select – rRNA HMR (Cat. No. / ID: 334385) was used for ribodepletion and paired end directional library was prepared using NEB NEBNext Ultra II Directional RNA Library Prep Kit for Illumina (E7760L), and run on Novaseq SP.

As mentioned above, data were pre-processed through sortMeRNA, trimmomatics. Rat transcriptome (Rnor_6.0.cdna+Rnor_6.0.ncrna) index was build and quantification was performed using Salmon^39^ (Quasi mode). Differential expression was analysed using DEseq2 package^40^. Criteria for differential expression: adjusted P value (Wald test followed by Benjamini-Hochberg FDR)<0.05 and |fold change|>2.

A list of differentially expressed genes and their normalized values can be found here: https://docs.google.com/spreadsheets/d/1IwR_XZgiLnHGcB7ubdInqGL8xdxM1nui/edit?usp=sharing&ouid=111216194520679821198&rtpof=true&sd=true

Off-target identification: A 125 bp region which is flanked by the qPCR primers to amplify RNLTR12-int (that region also includes our siRNA target) was used as an input in BLAT (using rat genome). Subsequently annotation was extracted for all the BLAT hits using HOMMER tools. A total of 19 genes were found whose intronic region contains a fragment of RNLTR12-int (either in the same or in the opposite strand), none were found in the exonic region. Differential expression (siRNLTR12-int vs control) of these 19 genes were extracted from the analysis above.

### Distribution of RNLTR12-int copies in rat genome

Repeatmasker annotation of repeats in the rat genome (rn6) were parsed through home written perl scripts. The following categories were used to identify a) RNLTR12-int sequence that is flanked by LTR: LTR to LTR total distance < 9 kb. At least 75% of the sequence flanked by the LTR should be repeatmasked. b) RNLTR12-int is not flanked by LTR: if RNLTR12-int fragmented sequences were separated by small insertions or other repeats up to 1 kb, they were assembled; and the whole contig size would be < 9 kb. In both (a) and (b): all features should be in the same chromosome and in same orientation.

### Identification of RNLTR12-int like sequence (RetroMyelin)

We employed the remote homology search algorithm nhmmer^41^, a probabilistic inference method based on a hidden Markov models, and used the RNLTR12-int consensus sequence (Repbase-GIRI) as an input to search for *RetroMyelin* in other genomes. The E-value threshold for inclusion was <0.01. Co-ordinates of the top hits were then queried using RepeatMasker^29^ to extract repeat annotation. Also, RepeatMasker^29^ was run (search engine: rmblast/hmmer/cross_match and selecting model organism as a DNA source) independently using the sequence of the top hits as an input to identify as well as to verify repeat type and family. Identified repeats which are just a simple low-complexity repeat (e.g. (TC)_n_) are not considered as *RetroMyelin* (Extended Data Table 1).

### Common Tree display

Using the taxonomic IDs of the subset of organisms as an input and by employing the Common Tree tool from NCBI, a hierarchical representation of the relationships among the taxa and their lineages was generated. The Phylip-format tree file was then visualized using iTOL^42^.

### Phylogenetic analysis

Sample collection: Top hit sequence of the identified *RetroMyelin* in different animals were used as an input for BLAT. For each BLAT result, the top 5 five highly similar sequences as well as 5 randomly selected sequences were collected. However, we do not consider sequences showing the following features in BLAT: a) the sequence alignment is such that only a small fragment of the full-length input sequence (e.g. 153 nt small fragment out of 1270 nt full length input) is aligned and or b) alignment is such that a small part is aligned to the genome followed by a long stretch of unaligned region, then another small part get aligned. Such sequences are so fragmented that they would not carry any functional copy. Furthermore, if sequences were not identified as ERV1-int by RepeatMasker^29^ run (search engine: rmblast/hmmer/cross_match and selecting model organism as a DNA source) then they were not considered further. In some animals we can not find 10 functional hits and therefore we collected samples from as many of them as were available.

Alignment and phylogenetic tree: MAFFT version 7 (a fast Fourier transform based multiple sequence alignment program)^43^ (keeping mafft flavour=auto) is employed to align *RetroMyelin* sequences. The alignment was curated using the BMGE (Block mapping and gathering with entropy) algorithm^44^. RAxML (RAxML-HPC BlackBox version 8.2.12)^45^ was employed for phylogenetic tree inference with maximum-likelihood and bootstrapping. Following the MRE-based Bootstopping criterion^46^. RAxML performed a rapid bootstrap search and collected 500 replicates. Trees were visualized in iTOL^42^.

### Rate heterogeneity analysis

Rate heterogeneity analysis was performed using BASEML (in PAML version 4.8a)^47^. This analysis is suitable for any sequences that have a common evolutionary origin^48^, and uses the phylogeny that describes their relationships in order to detect if there is variation in rates over sites where each site is assumed to have a constant rate over the tree, but potentially different from other sites. We used the REV+G (GTR+G) model with the assumption of the variation of evolutionary rates over sites following a gamma distribution. 4 gamma distributed rate categories were used. The parameters in the rate matrix (REV) were as described^49, 50^. The shape parameter (α) of the gamma distribution of rates over sites was estimated. H0 (no rate heterogeneity i.e., α→∞) was compared with H1 (gamma distributed rate heterogeneity) using a likelihood ratio test and the 2δ test statistics as described^23^. Following rejection of H0, site wise empirical Bayesian (posterior mean) relative rate estimates were computed as described^51^.

### EST mapping

Input nucleotide sequences were individually queried to expressed sequence tags (ESTs) database by using NCBI blastn suit. ESTs for which 0≤E-value≤6e-26 and 119≤alignment score≤2298, were mapped to the RNLTR12-int like sequences from the following animals: consensus sequence of Human (Dfam ID: DF0000205.5), mouse (Dfam ID: DF0001916.1), rat (RNLTR12-int from Repbase), zebrafish (Dfam ID: DF0002922.2). Consensus sequence was not available for elephant sharks, so we mapped to the genomic location obtained after nhmmer i.e. KI636671.1: 5058-5765 (Callorhinchus_milii-6.1.3).

### Animals

Sprague–Dawley rats were received from C. River (Margate, UK). Rats were maintained in individually vented cages at temperature 22°C ± 1°C and humidity 60% ± 5%, in a 12 hr light:dark cycle; food and water were supplied ad libitum in a standard facility for rodents at the University of Cambridge, UK. All animal studies were conducted under the Animals (Scientific Procedures) Act 1986 Amendment Regulations 2012 following ethical review by the University of Cambridge Animal Welfare and Ethical Review Body (AWERB).

### Isolation of OPC and OL

Postnatal day 7 (P7) rats were sacrificed using an overdose of pentobarbital and whole brain was dissected immediately in ice cold hibernate A low fluorescence (HALF) medium^52, 53^. Subsequently OPC was isolated as described earlier^52^. Per 10 million cell suspension, we used 2.5 μg mouse-anti-rat-A2B5-IgM antibody (Millipore; MAB312) and 20 μl of rat-anti-mouse-IgM antibody magnetic beads (Miltenyi, 130-047-302) for OPC isolation by magnetic-activated cell sorting (MACS). OLs were isolation from A2B5 negative fraction by using 2.5 μg of goat-anti-mouse-MOG-Biotinylated (RD systems, BAF2439) and 20μl mouse-anti-biotin magnetic micro beads (Miltenyi, 130-105-637).

Isolated OPC/OL were immediately placed in cell lysis buffer from mirVana (Ambion) RNA isolation kit. Otherwise, OPCs were seeded in 24-well-plate or 6-well-plate with poly-d-lysine (PDL) coated wells.

### Culture and differentiation of OPC

OPCs were maintained in OPC medium^52, 53^ [supplemented with 30 ng ml^−1^ basic fibroblast growth factor (bFGF) (Peprotech, 100-18B) and 30 ng ml^−1^ platelet-derived growth factor (PDGF) (Peprotech, 100-13A)] and 60μg ml^−1^ N-Acetyl cysteine (Sigma, A7250) in an incubator at 37°C and 5% CO_2_ and 5% O_2_. Medium was replaced in alternative days. OPC medium (100 ml) composition: A total volume of 100 ml of Dulbecco’s modified Eagle’s medium (DMEM)/F12 (Thermo Fisher, 11039-021) is supplemented with 1mM sodium pyruvate (Thermo Fisher, 11360-070), 5 mg apo-transferrin (Sigma, T2036), 1 ml SATO stock solution [described earlier (51)], 10μg ml^−1^ human recombinant insulin (GIBCO, 12585014).

For differentiation of OPC to OL: OPC medium was supplemented with 40 ng ml^−1^ 3,3ʹ,5-Triiodo-L-thyronine (T3) (Sigma; T6397)^52, 53^. Medium was replaced on alternate days for 5 days.

### Transfection

Lipofectamine™ RNAiMAX transfection reagent (ThermoFisher scientific) was used and followed the manufacturer protocol. 48 hours before transfection MACS isolated OPC (approx. 25,000) were seeded per well of a 24-well-plate (PDL coated) and cells were maintained in proliferation media. Before transfection media was replaced by differentiating media. 1 μl Lipofectamine™ RNAiMAX and 30pmol siRNA was used for transfection. siRNA against RNLTR12-int (Dharmacon) and control siRNA (siGENOME Non-Targeting siRNA pool #2) were procured from Dharmacon. 48 hours after transfection cells were harvested for RNA isolation. 5 days after transfection cells were fixed for immunofluorescence analyses.

siRNA sequence against RNLTR12-int: 5’ AAGUGAGGGCCUUCUAUGCUU 3’

### shmiR cloning and AAV PHP.eB packaging

SOX10 Multiple Species Conserved enhancer element 5 conjugated with cfos basal promoter (https://www.addgene.org/115783/)^54^ sequence was used for oligodendrocyte lineage specific expression of EGFP-shmiR with a bGHp(A) poly A signal (Extended Data Fig. 6a). This SOX10 driven entire sequence (Extended Data Fig. 6a) was cloned between the inverted terminal repeat of adeno associated virus 2 (AAV2ITR) vector. Cloning and large scale AAV PHP.eB packaging of the construct and purification was performed from AMS Biotechnology (Europe) Limited, UK. The viral tire was 10^13^ GCml^-1^. The following sequences were embedded into the miR-30 backbone of the shmiR vector:

shmiR-RNLTR12-int:

5’ ATAGAAGGCCCTCACTTTTA[GTGAAGCCACAGATG]TAAAAGTGAGGGCCTTCTATGC 3’

shmiR-Scrambled:

5’ TGGACGCTCGATTGATCATA[GTGAAGCCACAGATG]TATGATCAATCGAGCGTCCAAT 3’

The loop sequence is indicated by parenthesis.

### In vitro and in vivo infection

For *in vitro* study, OPC was isolated from rat pups using MACS (above) and approximately 30,000 cells were seeded to each well of a 24-well-plate, 48 hours before infection. Media was replaced to differentiating media (above) before infection. Approximately 5 × 10^10^ GC of virus was used per well. 4 days after infection total RNA was isolated using the mirVana kit (Ambion), DNase I (ThermoFisher scientific) treated before reverse transcription.

For *in vivo* experiment, 1μl of virus suspension (approximately 1 × 10^10^ GC) was injected stereotaxically into the cerebral cortex at 0.1mm anterior, 1.2mm lateral and 1.2mm deep with reference to Bregma under isoflurane anaesthesia, in neonatal rats at P1.

### Immunofluorescence staining

Cells were fixed with 4% PFA (10 min, room temperature), washed with PBS and proceeded for immunofluorescence staining as described^53^. For *in vivo* experiment, animals (at P14) were perfused using 4% PFA in PBS, brains were postfixed overnight with 4% PFA, cryoprotected with 20% sucrose, and embedded in OCT-medium (TissueTek). Immunofluorescence was performed on 12 μm cryostat section and closely followed the protocol as described earlier^55^. The following antibodies were used in this study:

| Primary antibody | Class | Host species | Dilution | Source | Catalogue number |
| --- | --- | --- | --- | --- | --- |
| Anti-MBP | IgG | Rat | 1 to 500 or 1 to 100 | Serotec | MCA4095 |
| Anti-OLIG2 | IgG | Rabbit | 1 to 1000 or 1 to 100 | Millipore | AB9610 |
| Anti-GFAP | IgY | Chicken | 1 to 1000 | Abcam | ab4674 |
| Anti-O4 | IgM | Mouse | 1 to 1000 | R&D Systems | MAB1326 |
| Anti-CC1 | IgG2b | Mouse | 1 to 100 | Merck/milipore | OP80 |
| Anti-GFP | IgY | Chicken | 1 to 100 | Abcam | ab13970 |
| Alexa Fluor 488 Anti-Mouse | IgM | Goat | 1 to 1000 | Invitrogen | A21042 |
| Cy<sup>TM</sup>3 AffiniPure Anti-Rat | IgG (H+L) | Donkey | 1 to 1000 or 1 to 500 | Jackson Immuno Research | 712-165-150 |
| Alexa Fluor 647 Anti-Rabbit | IgG | Donkey | 1 to 1000 or 1 to 500 | Invitrogen | A31573 |
| Alexa Fluor 488 Anti-Chicken | IgG (H+L) | Goat | 1 to 1000 or 1 to 500 | Invitrogen | A11039 |
| Alexa Fluor-568-Anti-mouse | IgG2b | Goat | 1 to 500 | Invitrogen | A21144 |

Images were acquired with an SP5 (Leica TCS) or SP8 (Leica TCS) confocal microscope. All acquisition settings were kept constant for the whole experiment. Images were processed using the ImageJ version 2.1.0 software^56^. For *in vivo* immunoassayed sections, integrated density of MBP immunostaining was measured using ImageJ 2.1.0. Cell counting was performed using Cell Counter tool in ImageJ 2.1.0. Immunofluorescence experiments were performed and analysed blind to treatment.

### qRT-PCR analysis

Total RNA was isolated using the mirVana kit (Ambion). Total RNA was DNase I (ThermoFisher scientific) treated before reverse transcription. cDNA was generated by using random hexamer (Invitrogen) and SuperScript™ IV Reverse Transcriptase (Invitrogen™) following manufacturer protocol. Real time PCR was performed using SYBR green master mix (Applied Biosystems, ABI) in QuantStudio 7 Flex (Applied Biosystems, ABI) machine. The real time PCR program was 50°C/ 2 min, 95°C / 10 min., 40 cycles 95°C/ 15 s, 60°C/ 1min. Dissociation protocol was run.

Rat specific primer sequences used in qPCR:

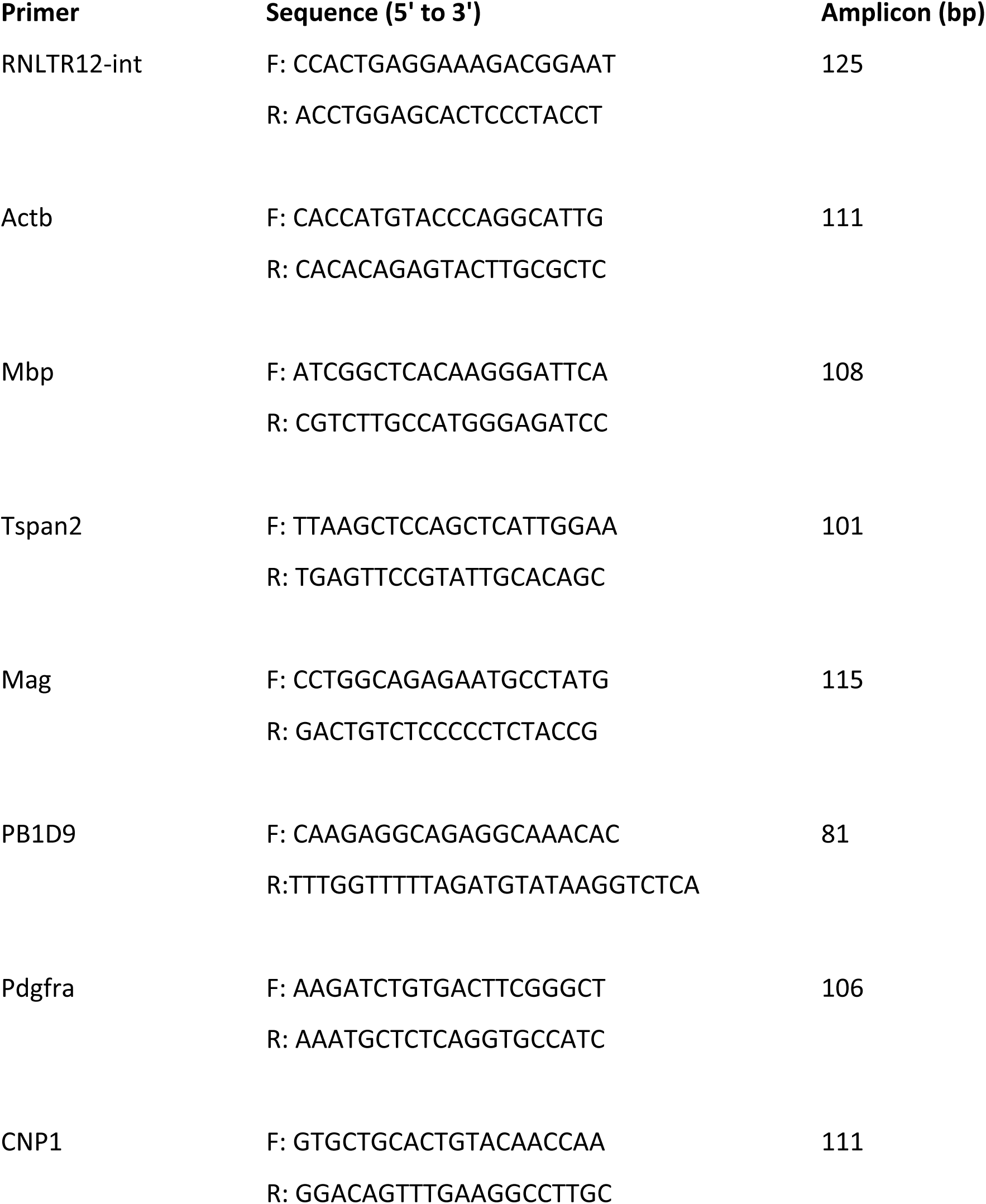

RNLTR12-int sequence was located in multiple chromosomes which can be identified using Affymetrix probe set (1379497_at). We did BLAT (from UCSC) and designed the primers from the top hit (chr7:82148213-82148189, assembly: rn5), which was one among many that showed 100% identity. Since RNLTR12-int is intron-less, we excluded the possibility of trace genomic DNA (gDNA) in our RNA samples by treating with DNase I and designing intron flanking primers targeting cytoplasmic beta actin (Actb) gene to detect gDNA contamination. We observed only cDNA specific expected size amplicon band (approx. 125 bp) in our OPC and OL samples (Extended Data Fig. 2a). Further melting curve analysis by quantitative PCR (qPCR) yielded only single peak, suggesting a single amplicon (Extended Data Fig. 2a), confirming that there was no gDNA contamination.

Actb was used as an endogenous gene for normalization control. We used Actb as a normalization control because in our microarray data analysis there was no expression difference of this gene between OPCs and OLs was observed (Extended Data Fig. 2b). Relative quantification of qPCR data was analyzed by Paffl method^57^. PCR efficiency of the target gene and the endogenous control was determined from the slope of the respective standard curve.

### RNA immunoprecipitation (RIP)

Crosslinking RNA immunoprecipitation was performed by following standard Abcam protocol. 5-6 Sprague–Dawley rat brains (P7) were used. Meninges were removed and cellular dissociation^52^ was performed up to 33% Percoll step to remove debris and fat. Approximately 4-6 X 10^7^ cells were obtained, re-suspended in Hank’s balanced salt solution (HBSS) and immediately cross-linked with formaldehyde (with a final concentration 1%) for 10 min at room temperature. Crosslinking was stopped with glycine (final concentration 0.125M) and subsequently washed with PBS. Nuclei was isolated in nuclei isolation buffer and nuclear pellet was re-suspended in RIP buffer. Sonication (30 s on, 30 s off, 10 min) was performed using Bioruptor (Diagenode). Pre-cleaning of lysate was carried out using blocked Protein G sepharose (ab193259, Abcam). Blocking of Protein G sepharose was carried out by using 500ng/ul yeast tRNA and 1mg/ml RNase-free BSA. A total of 45% (450 μl) of pre-cleaned lysate was used for anti-SOX10 (sc-365692, Santa Cruz) or anti-immunoglobulin G1 (IgG1) (sc-3877, Santa Cruz) samples, and 10% was used for input sample. A total of 4 ug antibody (SOX10 or immunoglobulin G1) was used for immunoprecipitation, and blocked Protein G sepharose was used for the pull-down step after immunoprecipitation. Eluted solution was de-crosslinked and proteinase K treated, DNAase treatment (TurboDNAse) was performed and RNA was isolated. RNA samples were analyzed by qPCR.

### Chromatin-immuno-precipitation (ChIP)

Four P7 rat brains were pooled each time for OPC isolation and approximately 2×10^5^ cells were seeded per well of a six-well-plate (PDL coated). Transfection were performed as mentioned above. 4 days after transfection, cells were cross linked with 16% HCHO added per well onto the media (final concentration 1%) and kept for 10 min and subsequently followed the standard ChIP protocol^58^. After fixation and quenching by glycine (final concentration: 0.125 M), cells were dissociated from PDL-coated wells using TrypLE Express (Thermo Fisher; 12604013).

Sonication (30 s on, 30 s off, 30 min)^59^ was performed using a bioruptor (Diagenode) and the fragment size 200–500 bp was verified. Protein G Sepharose beads were blocked using 1 mg/ml BSA (stock 10 mg/ml). As described in RIP method (above), lysate was pre-cleaned and immunoprecipitation using antibody (anti SOX10 or IgG1) and blocked Protein G sepharose was used for the pull-down step after immunoprecipitation. Washing and preparation of DNA was performed as described^58^. Precipitated DNA samples were analysed by PCR and qPCR. The following primers were used for PCR after ChIP:

Mbp promoter:

FP: 5’ CATTGTTGTTGCAGGGGAGG 3’

RP: 5’ GCTCGTCGGACTCTGAGG 3’

### Cloning, in vitro transcription (IVT) and labelling of RNA

Consensus sequence of RNLTR12-int (obtained from Repbase-Giri: https://www.girinst.org/repbase/) was cloned in pcDNA3.1 (+) using the restriction site NheI/XbaI.

Purified linearized plasmid (from above) was used as a template and transcribed *in vitro* using HiScribe™ T7 ARCA mRNA Kit (with tailing) (New England Biolabs) following manufacturer protocol (E2060) and purified using Direct-zol^TM^ RNA Miniprep kit (Zymo Research).

For surface plasmon resonance control, the following RNA was synthesized from Dharmacon: 5’ CCUGAUUUUUAAGGAAUAUCGCAAGAAUGCCGCGAAUGAAAAA 3’

RNA was 3’ labelled with biotin using Pierce™ RNA 3’ End Biotinylation Kit (ThermoFisher) and purified using Monarch® RNA Cleanup Kit (New England Biolabs).

### Surface plasmon resonance (SPR) experiment

The SPR experiment was performed on a Biacore T200 (GE Healthcare/Cytiva) using a Series S Sensor Chip SA (Cytiva) following the manufacturer’s instructions. Tris Buffered Saline (TBS) pH7.4, supplemented with 0.002% Tween20, 100 mM glycerol and 5% glycine, was used as the running buffer throughout. Immobilization was achieved by flowing approximately 5 ugml^-1^ biotinylated RNA for 300 seconds at a flow rate of 2 ulmin^-1^. Recombinant Human SOX10 protein (Euprotein) at a concentration of 2 uM was then flowed as analyte for 60 seconds at a flow rate of 30 ulmin^-1^. Data were analysed using the inbuilt BIAEval software (Cytiva).

### Zebrafish experiment

Zebrafish were maintained in the University of Edinburgh BVS Aquatics facility under project license PP5258250. To induce mutations in *RetroMyelin* loci in the zebrafish genome, we designed a guide crRNA targeting the sequence TAATGAAGCAATCAAACAAG*TGG* (PAM sequence italicized), which is conserved in the top 10 hit loci most similar to the RNLTR12-int. Ribonucleoprotein complex (RNP) were prepared by annealing 20 µM crRNA with 20 µM tracrRNA (IDT DNA) at 95°C, and incubating 1.6 µM of annealed gRNA with 2 µM Engen Cas9 enzyme (New England Biolabs) at 37°C for 10min. 1nL RNP or 1nL of Cas9-only control solution were microinjected into fertilized one-cell stage eggs of the transgenic myelin reporter line Tg(mbp:EGFP-CAAX)ue2^60^. Embryos were kept at 28.5°C in 10mM HEPES-buffered E3 Embryo medium.

At 5dpf, larvae were anesthetized in 0.16mg/mL tricaine (ethyl 3-aminobenzoate methanesulfonate salt, Sigma-Aldrich) and loaded and oriented for imaging using an LPSampler and VAST BioImager fitted with a 600 μm capillary (Union Biometrica) as previously described^61^. We imaged the whole length of the embryo with a Zeiss Axio Examiner D1 equipped with a CSU-X1 spinning confocal scanner, a Zeiss AxioCam 506 m CCD camera and a Zeiss C-Plan-Apochromat 10x 0.5NA objective, by tiling 5 z-stacks per embryo with a z-step of 3 μm. This system automatically collects individual larva from a multi-well plate, positions every larva consistently in the same angle, acquires a brightfield image for morphological analyses, and a high-resolution optically-sectioned GFP image for myelination analyses. Subsequently, individual larva was dispensed into a multi-well plate, from which we collected genomic DNA using the HOTSHOT method^62^.

For analyses of RNP activity, we amplified a 607 bp PCR product around the target gRNA sequence from genomic DNA of imaged larva using primers 5’-TTC GGC TGC GGT AGC AAA-3’ and 5’-CGC TGT CTC AGT GGT CAG-3’, and Onetaq DNA polymerase (New England Biolabs) according to the manufacturer’s instructions (briefly, 94°C-30s, 35X [94°C-30s, 54°C-30s, 68°C-50s], 68°C-2min). PCR reactions were run in a 1% agarose gel, and a single band of the expected size was purified using Monarch Gel Extraction Kit (New England Biolabs). Purified amplicons from 6 Cas9-only control injected larva and from 24 gRNA-injected (and imaged) larva were sent for Sanger sequencing using the PCR primers (Source Bioscience).

For morphology analyses of brightfield images, we used Fiji/ImageJ to mark the rostral and caudal ends of each larva (mouth and tail fin) to measure body length, and the ‘wand’ tool (with a constant threshold set manually with the first 2-3 samples and used throughout the dataset) to select the eye area while blinded to experimental condition. For myelination image processing and analyses, maximum intensity projections were created for all samples and a semi-automated Fiji/ImageJ script defined regions of interest in spinal cord (whole, dorsal and ventral as well as bins along the anterior-posterior length) in a manner blinded to experimental group^61^. The mean gray value was obtained for each region of interest. In the figures, all zebrafish images represent a lateral view, anterior to the left and dorsal on top. For the myelination figure panel, control and gRNA-injected images were prepared in parallel using the same brightness and contrast settings, and the ‘fire’ look-up table was used to convey fluorescence intensity/gray values as colour changes, indicated in the calibration bar.

### Frog (Xenopus laevis) experiment

Xenopus were raised in the animal facility of Institut du Cerveau – ICM, Paris, France (agreement # A75-13-19) under the project license approved by the ethical committee of the French Ministry of Higher Education and Research (APAFIS#5842-2016101312021965). Maintenance of tadpoles, Crispr/Cas9 injection, albino selection and immunolabeling procedures were performed as described in Mannioui et al.^63^. Preparation of sgRNAs, generation of S1P5 null mutant: CRISPR sgRNA were as described^63^. We design three guide RNAs (gRNA) using CHOPCHOP (https://chopchop.cbu.uib.no/) with 1 to 3 mismatches capable of covering a target region in multiple sequences of *Xenopus RetroMyelin*:

(The red color indicates the mismatch in gRNA, the yellow color indicates the PAM sequence.)

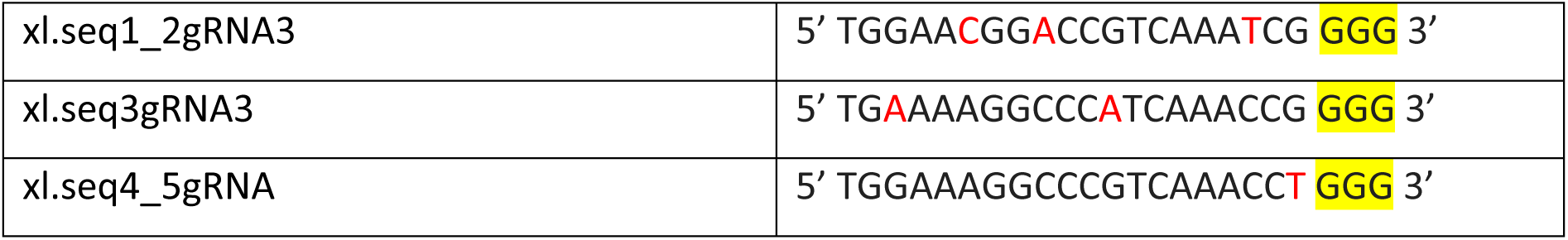

The region including the guide RNA target site were amplified by PCR using primers designed with Primer3, and Sanger sequencing with the forward primers was performed. The sequences data were analyzed with CRISP-ID^64^.

FP: 5’ GTGCATGGTCAGGTGTTCTC 3’

RP: 5’ GGATCCACGCATCCCACTGCAA 3’

Images were acquired using a Nikon A1R-HD25-confocal microscope. Z-series of wholemount tadpole were performed at 0.375 μm step. Maximum orthogonal projection of images was carried out using Fiji software (NIH, Bethesda, Maryland). To analyze of Olig (this is not OLIG1 or OLIG2, for simplicity reason we wrote XMM)^65^ and MBP expression, images were quantified for labeled area fraction (%) by automated counting using ImageJ software. Briefly, images were normalized by subtracting background and a threshold (MaxEntropy) was applied. Mean values of area fraction (%) were obtained from images of tadpole injected with gRNA vs control.

### Statistical analysis

R (version 4.0.2) (http://www.R-project.org) or GraphPad Prism (version 8.4.3) was used for statistical analysis. Shapiro-Wilk test^66^ was performed to test normality. Welch’s t test was performed whenever heteroscedasticity existed in the data set.

---

**Extended data Fig. 1.**
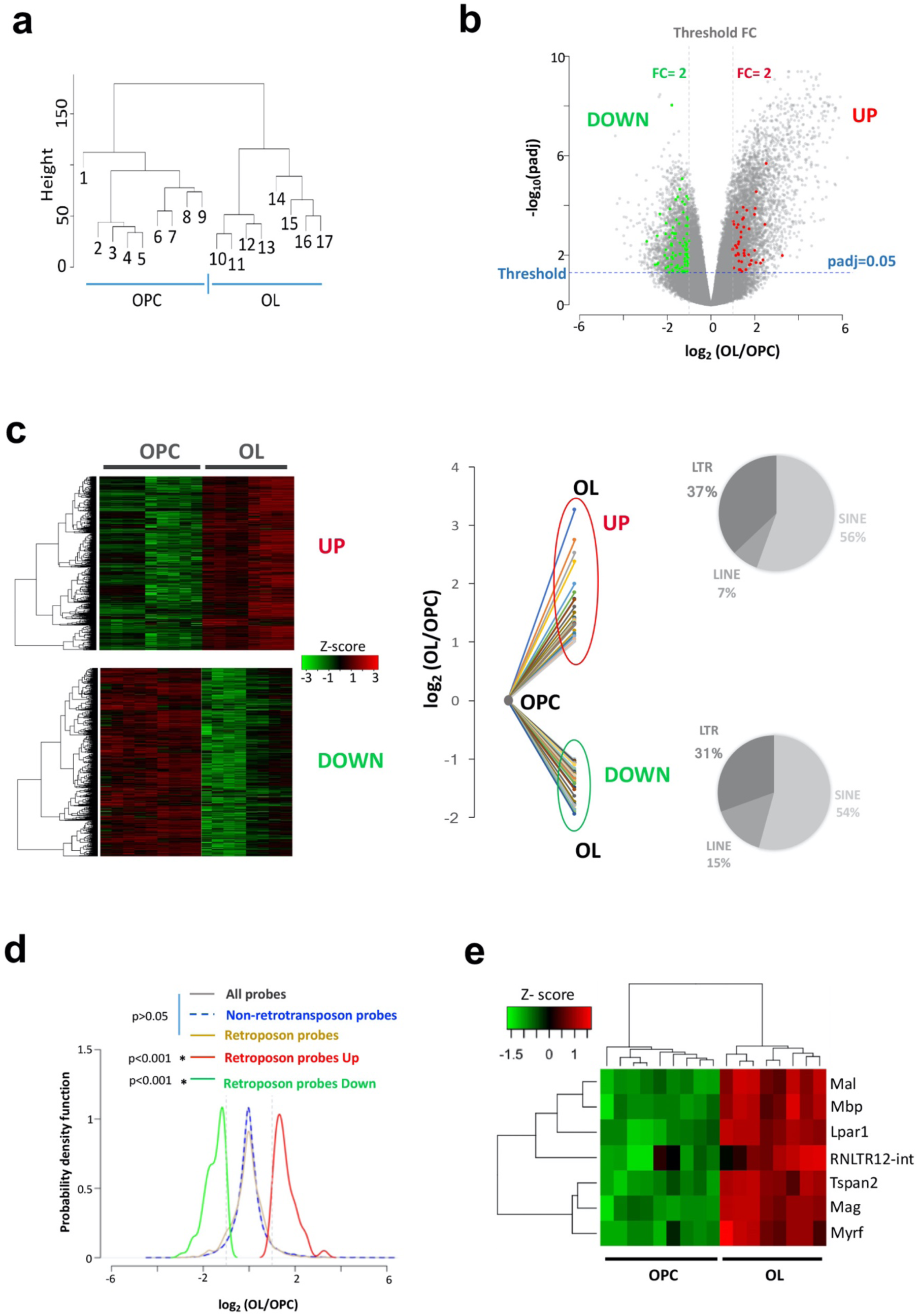
cGenome wide expression of retrotransposons in OPC and OL. **a,** Dendrogram representation of OPC and OL samples, showing OPC and OL samples were formed two separate clusters. Average linkage hierarchical clustering (using Euclidean distance matrix) was performed with respect to the intensity levels of probes in Affymetrix GeneChip. **b**, Probe level analysis of differential expression. Differentially expressed retrotransposons probes (red and green dots) and probes for protein coding genes (grey dots). Probes for which the expression difference between OL and OPC is 2-fold up or down and the adjusted p value<0.05 were considered differentially expressed. FC: Fold change. Padj: adjusted p value (Benjamini Hochberg FDR). **c**, Left, Heatmap plot and dendrogram representation of all differentially expressed probes in OL and OPC samples. Right, Differential gene expression (log_2_ value) of retrotransposon probes were plotted. UP: induced probes, DOWN: repressed probes. Proportion of different retrotransposon type in induced and repressed probes were plotted in a pie chart. **d**, Probability density plots showing the behavior of retrotransposon probes, non-retrotransposon probes, all probes and differentially expressed retrotransposon probes. Vertical dotted lines: 2-fold change of expression. * p<0.05, Wilcoxon rank sum test. **e**, Expression levels of RNLTR12-int and myelination hubs were represented as heat map and a dendrogram derived after two-way hierarchical clustering.

**Extended Data Fig. 2.**
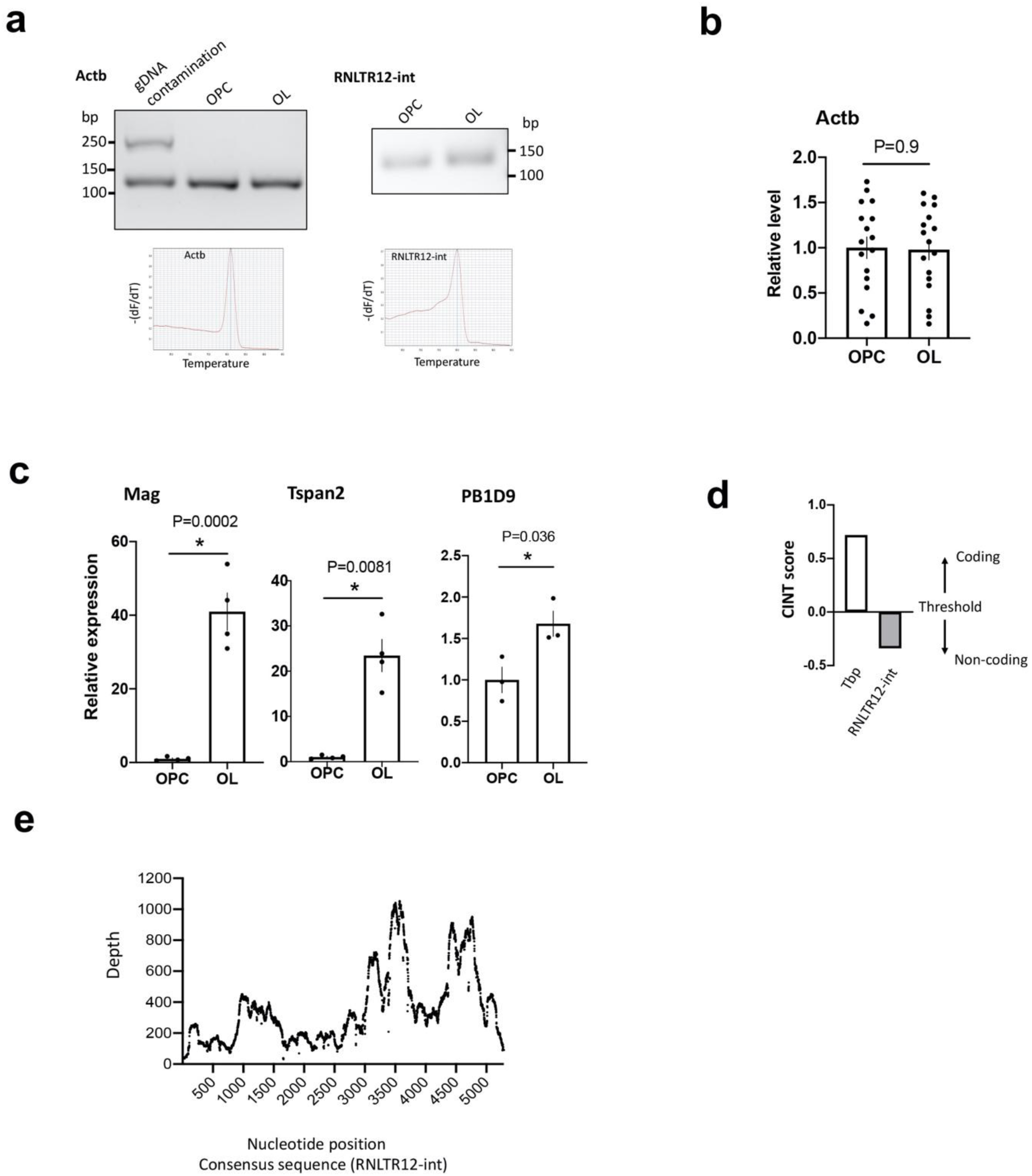
Validation of RNLTR12-int expression in MACS isolated OPC (A2B5+) and OL (Mog+). **a**, Top: (left) RT-PCR analysis of cytoplasmic beta-actin (Actb) using intron flanking primers. gDNA contamination in RNA resulted an upper band. Only single lower band obtained in OPC and OL samples, confirms no gDNA contamination. (right) RT-PCR analysis of RNLTR12-int expression in OPC and OL. PCR product was run on a 2% agarose gel. Bottom: qPCR melting curve analysis resulted only single peak for Actb (left) and RNLTR12-int (right). **b**, Actb expressions remained unchanged in OPC and OL as determined by Affymetrix GeneChip. Normalized intensity of all Actb specific probes (17 probes) were used and plotted relative to OPC. N=9 (OPC), 8 (OL), p>0.05, Student’s t-test (unpaired, two-tailed). mean±SEM. **c**, Expression levels of Mag, Tspan2 and PB1D9 is elevated in OL as compared to OPC determined by RT-qPCR. N=3-4, *p<0.05, Student’s t-test (unpaired, two-tailed), mean±SEM. **d**, RNLTR12-int encoded transcript is likely to be a non-coding RNA as predicted by CNIT algorithm^34^. Score of a known protein coding gene, TATA-box binding protein (Tbp, Transcript ID: ENSRNOT00000002038.4) is shown. **e**, RNA sequencing reads were aligned to RNLTR12-int consensus sequence and the depth was plotted.

**Extended Data Fig. 3.**
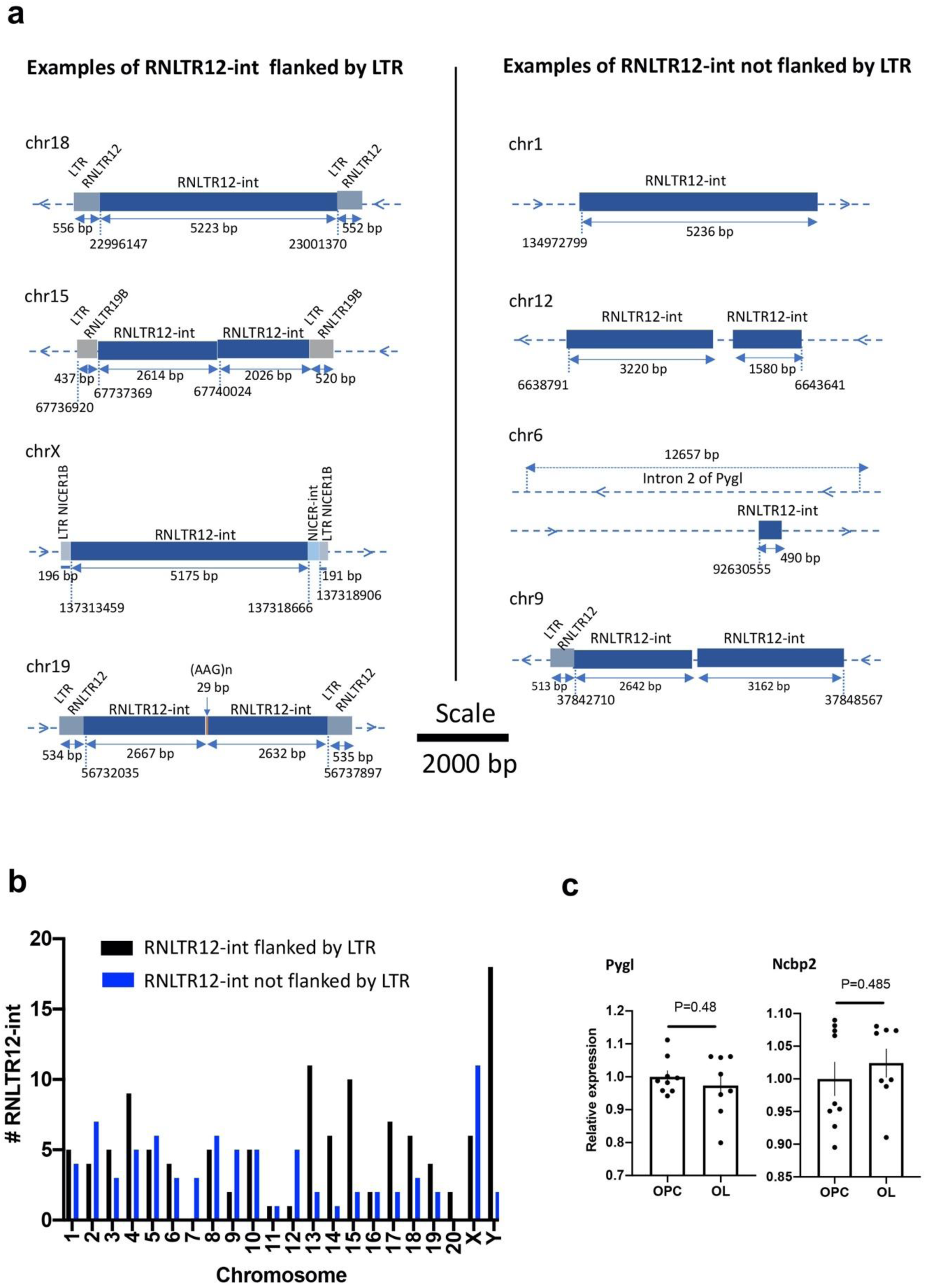
Schematic view of some examples of RNLTR12-int in rat genome (rn6) which are flanked (**a**, left) or not flanked (**a**, right) by long terminal repeat (LTR). length in base pair (bp) is written in the middle of the double headed arrow. Genomic location is mentioned below of a vertical dotted line. On each horizontal line the blue colored symbol ‘>’ indicates forward strand, ‘<’ indicates reverse strand. chr: Chromosome. **b,** Total number of RNLTR12-int (either flanked or not flanked by LTR) per chromosome in rats which were identified by our criteria (see methods) is plotted as a bar diagram. **c**, Expression levels of Pygl and Ncbp2 in OPC and OL (determined by Affymetrix GeneChip) were plotted relative to OPC samples. N=9 (OPC), 8 (OL), Mean±sem, p>0.05, Student’s t-test (unpaired, two-tailed).

**Extended Data Fig. 4.**
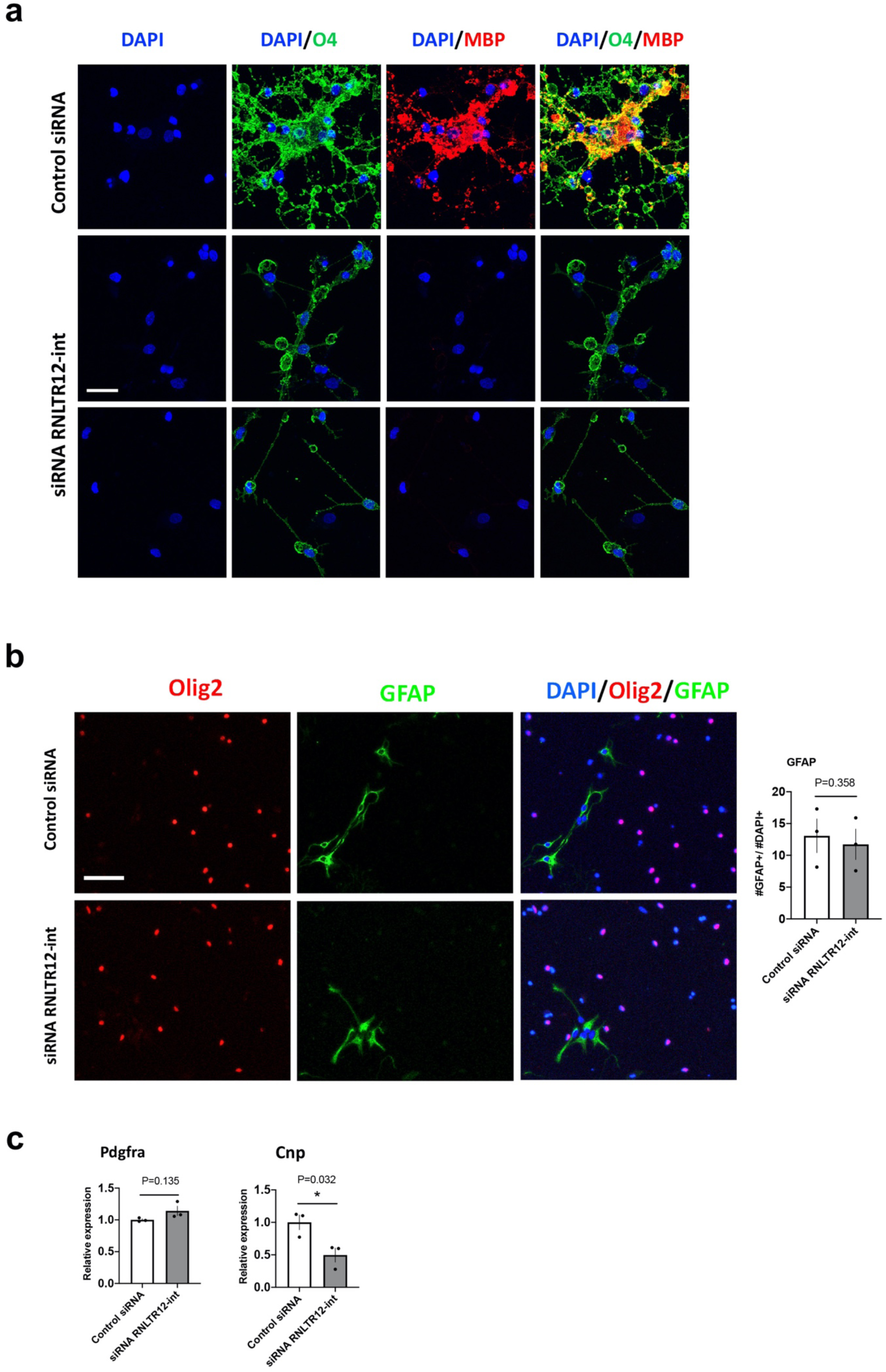
**a**, Absence of complex oligodendrocyte morphology due to inhibition of RNLTR12-int. Immunofluorescence analysis O4 immunostaining 5 days after transfection of siRNA. Scale bar: 26 μm. siRNLTR12-int: siRNA against RNLTR12-int, control siRNA: siGENOME non-targeting siRNA pool (Dharmacon). Representative image of 3 independent experiments. **b,** Inhibition of RNLTR12-int did not initiate astrocytic fate. Immunofluorescence analysis of GFAP immunostaining 5 days after transfection of siRNA. Scale bar: 60μm. Percentage of GFAP+ cell in total DAPI+ cells were plotted. N=3 independent experiments (each time 3 replicates), mean±SEM, p>0.05, Two-way ANOVA. siRNLTR12-int: siRNA against RNLTR12-int, control siRNA: siGENOME non-targeting siRNA pool (Dharmacon). **c**, Effect of the inhibition of RNLTR12-int on RNA level expression of Pdgfa and Cnp were determined by RT-qPCR. Data were normalised to Actb. N=3 independent experiments, mean±SEM, * p<0.05, Student’s t test (unpaired, two tailed).

**Extended Data Fig. 5.**
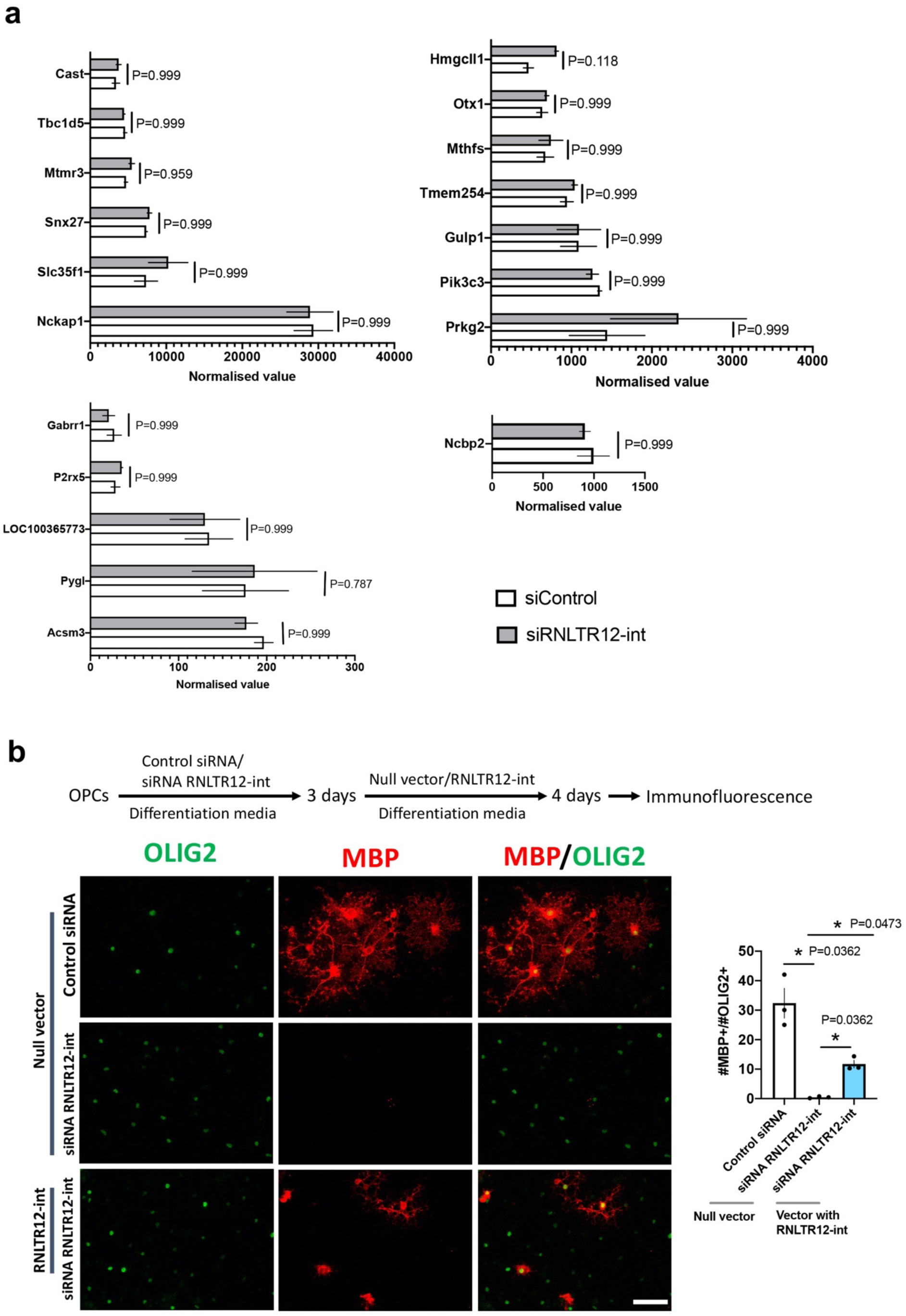
**a**, Exclusion of any off-target effect. siRNA-mediated inhibition of RNLTR12-int in differentiating OPC does not alter expression of genes whose intronic region (either in the same or the opposite strand) contains fragment of RNLTR12-int. Normalized values from RNAseq data plotted, mean±sem, P>0.05, Benjamini-Hochberg FDR after Wald test. **b**, Overexpression of RNLTR12-int elevates MBP expression in differentiating OPCs. pcDNA 3.1 (+) bearing a full length RNLTR12-int consensus sequence or null vector was transfected 3 days after siRNA administration (schema above). Representative immune-stained image (left) and quantification of MBP+ oligodendrocytes (right) shown. * adjusted p < 0.05, Brown-Forsythe and Welch’s ANOVA (Benjamini-Hochberg FDR post test), mean+SEM, N=3 independent experiments.

**Extended Data Fig. 6.**
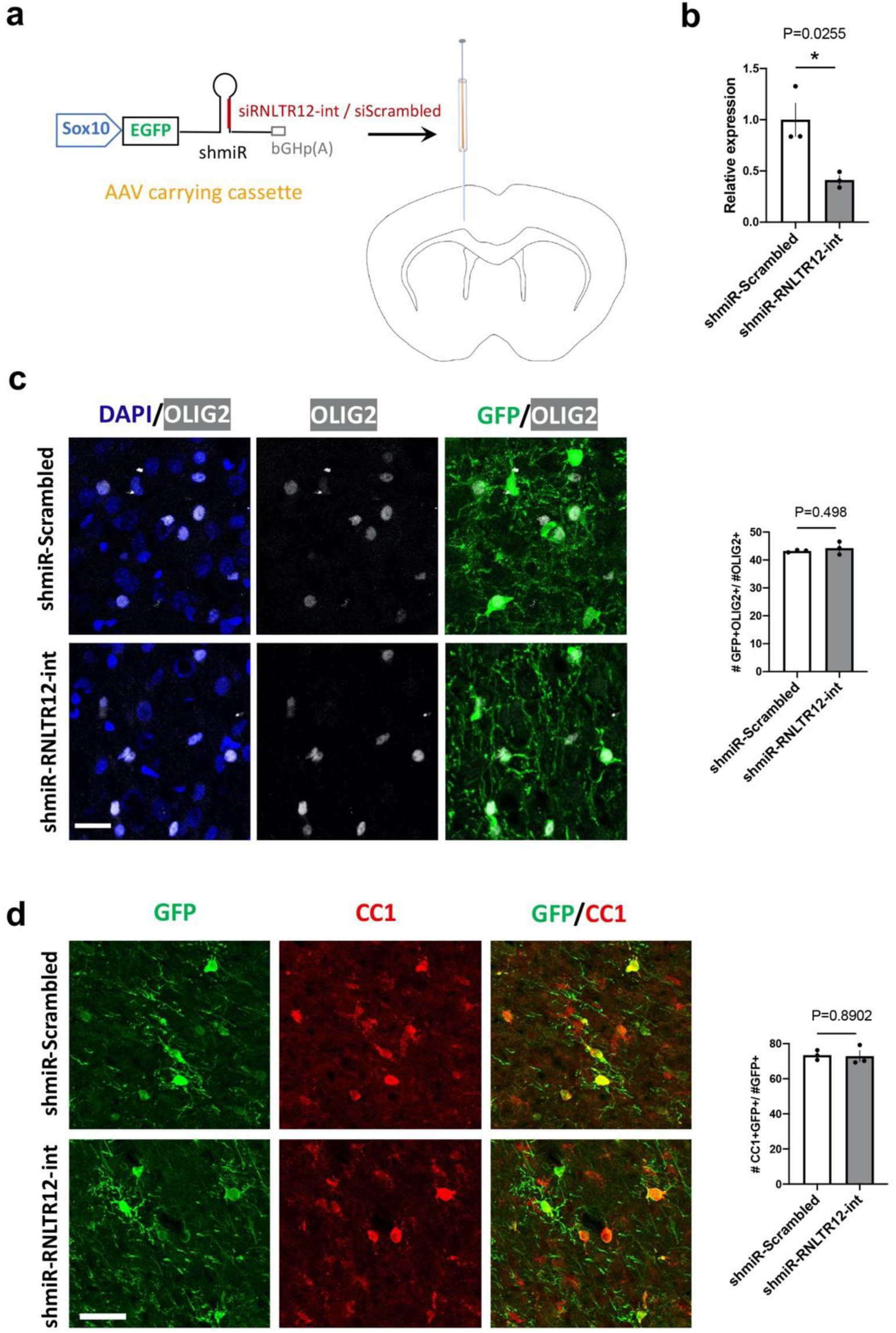
**a**, SOX10 driven EGFP (Emerald GFP) construct that carry shmiR where shRNA is embedded into a microRNA (miR-30) cassette. bGHp(A): Bovine growth hormone poly A signal. This construct is SOX10 driven and expresses EGFP. This also produces shRNA that function through RNAi pathway. Shown is the rat brain area where AAV (carrying SOX10-EGFP-shmiR-bGHp(A) were injected. **b**, AAV carrying the above construct were infected into the cultured OPC and allowed to differentiate for 4 days, then RNA was isolated. RT-qPCR analysis of RNLTR12-int revealed its reduced expression in shmiR-RNLTR12-int infected cells. N=3 independent experiments, mean+SEM, * p<0.05, Student’s t test (unpaired, two tailed). **c-d**, AAV carrying the above construct were injected into newborn rat brain (at P1). Brains were harvested at P14 for immunofluorescence. **c**, Left: immunofluorescence analysis of OLIG2 and GFP immunostaining. Right: quantification of OLIG2+GFP+ cells and plotted as a percentage to OLIG2+ cells. N=3 rats, mean+SEM, p>0.05, Student’s t test (unpaired, two tailed). **d**, Left, immunofluorescence analysis of CC1 and GFP immunostaining. Right, quantification of CC1+GFP+ cells among GFP+ cells and represented as a percentage. N=3 rats, mean+SEM, p>0.05, Student’s t test (unpaired, two tailed).

**Extended Data Fig. 7.**
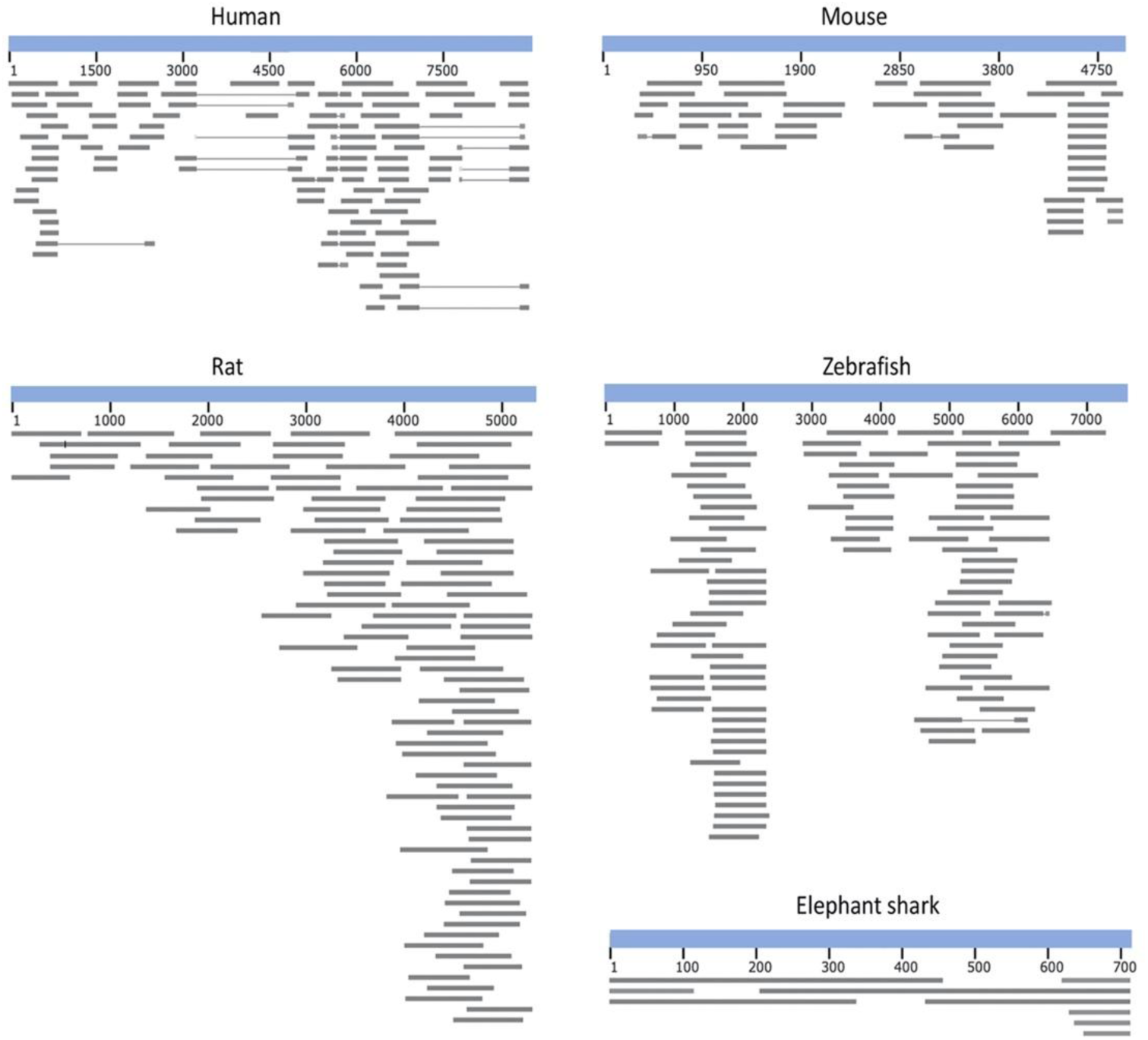
Expressed sequence tags (ESTs) were aligned to the RetroMyelin sequence of human, mouse, rats, zebrafish and elephant sharks. BLASTN was used to search ESTs. For the displayed ESTs: 0<E-value<6e-26 and 119<alignment score<2298. For human, mouse, rat and zebrafish: consensus sequence was used. For elephant shark: KI636671.1: 5058-5765 (Callorhinchus_milii-6.1.3) was used (this was the best hit obtained after nhmmer search on RNLTR12-int (see methods); E-value=1.7e-14).

**Extended Data Fig. 8.**
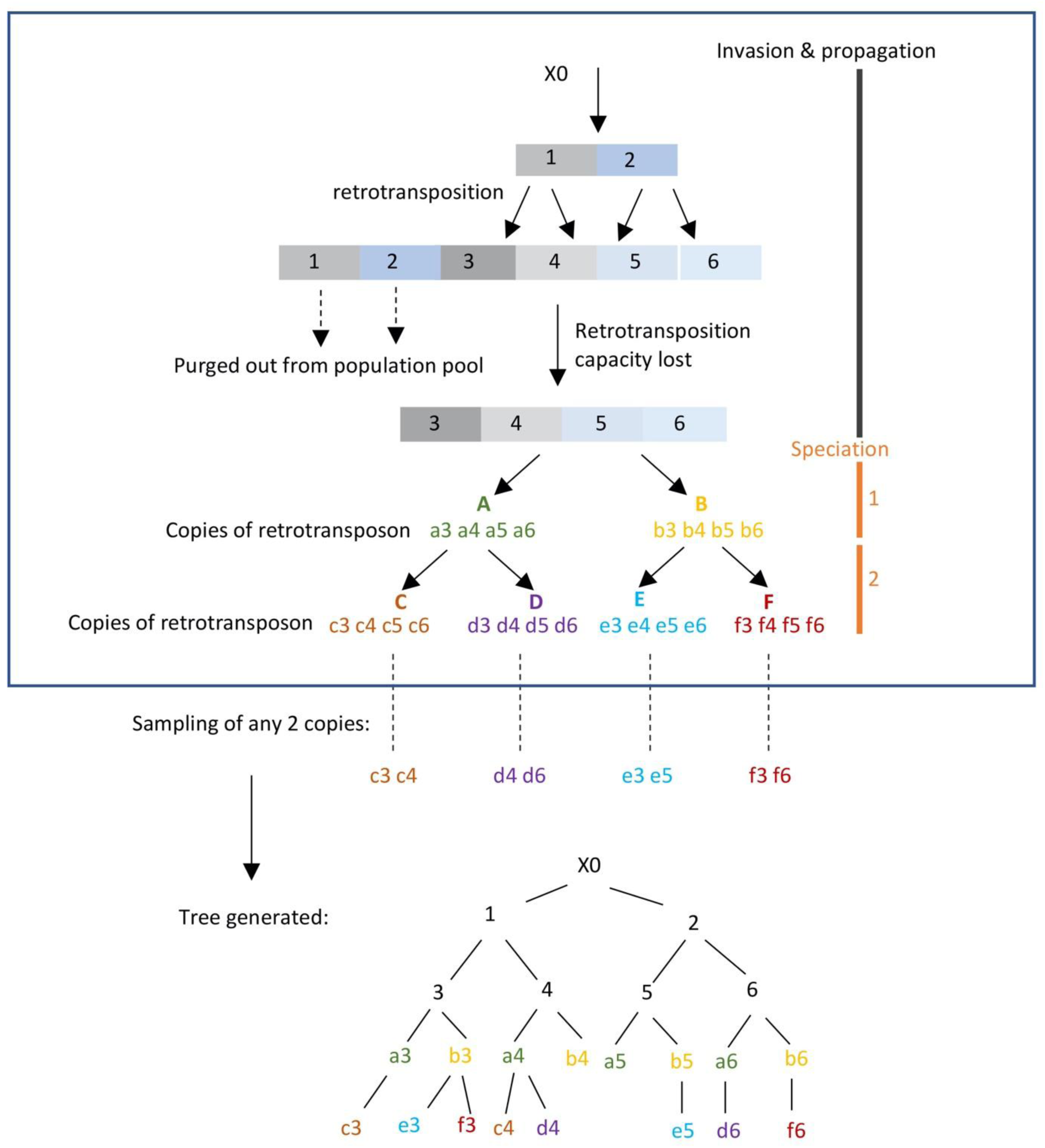
A representation of a hypothetical phylogenetic tree reconstructed from copies of a specific retrotransposon sequence existing in different species. Let us assume that a viral invasion had happened before speciation. The newly invaded proviral genome (X0) generates copies 1 and 2 in a host genome. Subsequently retrotransposition of copy 1 propagated as copies 3 & 4, while copies 5 & 6 were generated from copy 2. 3, 4, 5 and 6 completely lose their capacity to further retrotranspose due to mutations associated with each recombination event. Selective forces acting at the level of host population subsequently purge copies 1 and 2 from the population pool. Speciation then produces species A and B, and from these ancestors C & D and E & F, respectively, are generated (speciation 2). Assume that each species received the versions of the original copy of 3, 4, 5, 6 (represented by a, b, c, d, e, f). A scientist collects samples of two copies (randomly) from each extant species (C, D, E, F) for reconstruction of a phylogenetic tree. In the resulting tree, we can see that the phylogeny of the retrotransposon copies (c3, e3, f3, c4, d4, etc.) does not recapitulate the species tree and the copies sampled from each species are not clustered together.

**Extended Data Fig. 9.**
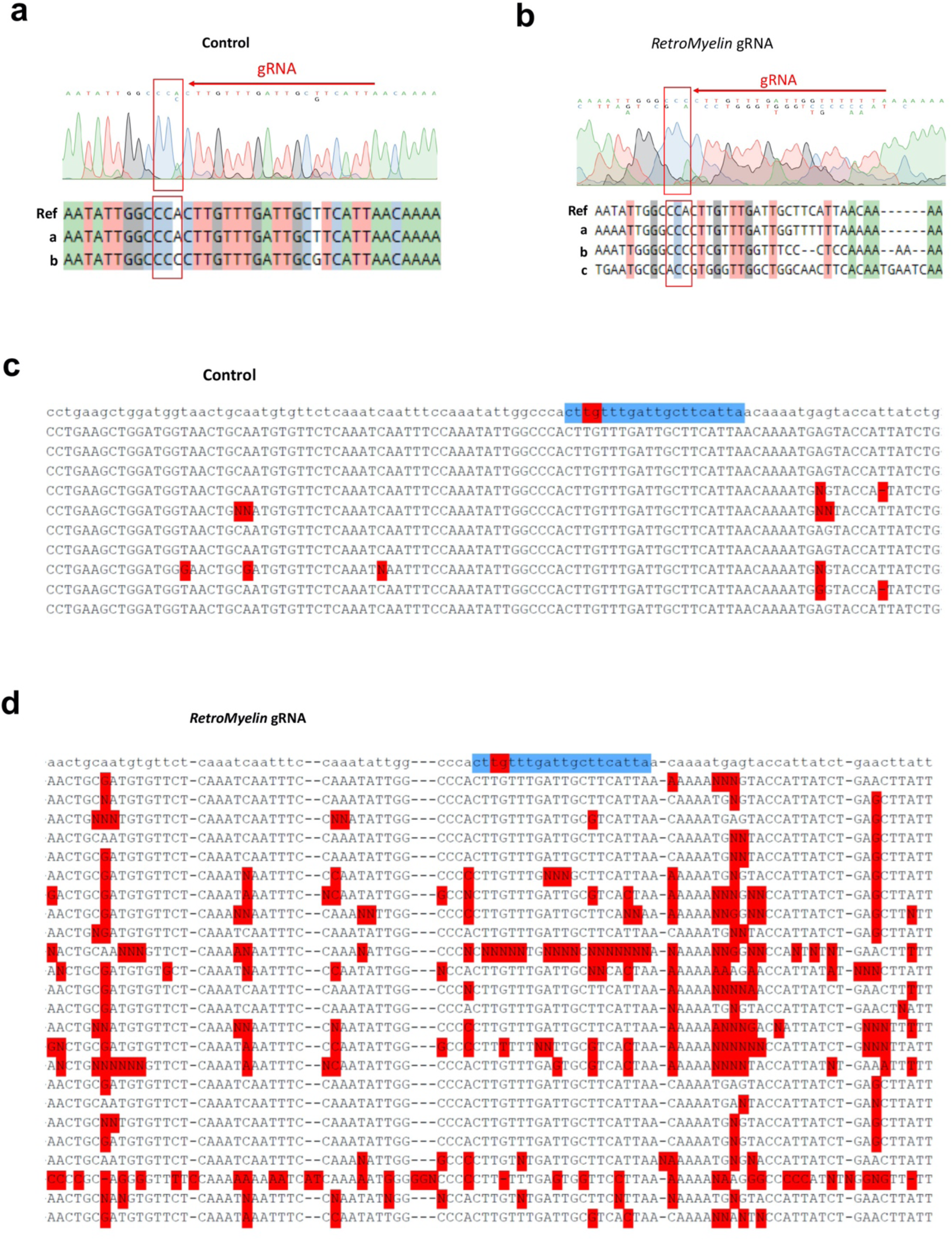
**a-b**, Example chromatograms from Sanger sequencing of a PCR product amplified from genomic DNA extracted from a control-injected larva (**a**) or *RetroMyelin* gRNA-injected larva (**b**). Predicted genomic sequence indicated on top, and gRNA target and PAM sequence indicated by red box and arrow. Below the traces are predicted alleles identified from the chromatograms using CRISPR-ID^64^, showing only few SNPs in control, but multiple indels and variations in the gRNA-injected larva. **c-d**, Alignments of multiple sequencing results for PCR products amplified from genomic DNA extracted from control (**c**) or *RetroMyelin* gRNA-injected (**d**) larva to the reference sequence (top, gRNA sequence in blue and predicted cut site in red). Note widespread mismatches in *RetroMyelin* gRNA-injected traces.

**Extended Data Fig. 10.**
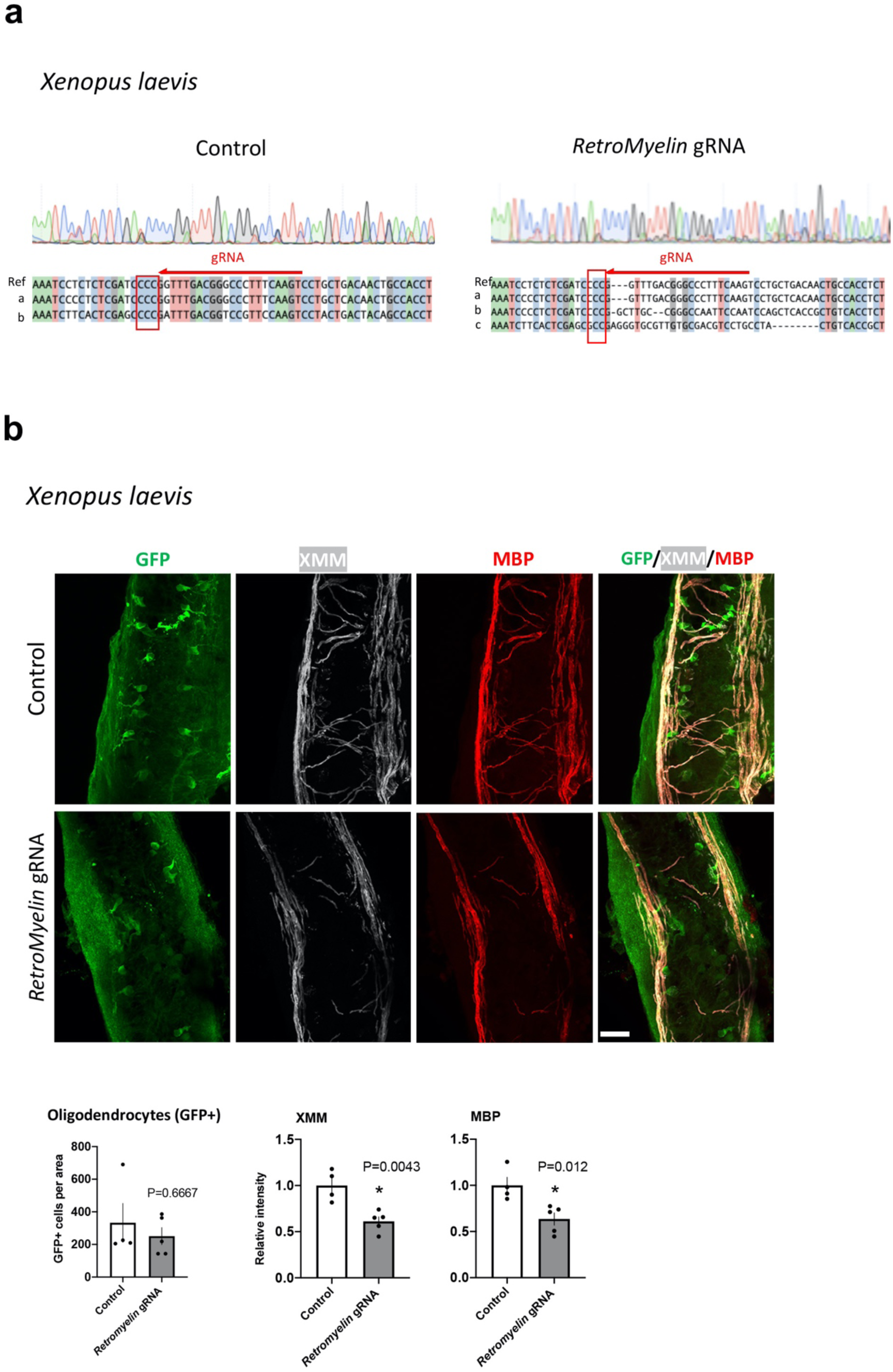
**a**, Analysis of overlapping peaks arising from control (left) or *RetroMyelin* gRNAs-injected Xenopus (right) using CRISP-ID^64^. Shown is representative chromatogram from Sanger sequencing of PCR amplicons from genomic DNA extracted from control (left) or *RetroMyelin* gRNA-injected *Xenopus* tadpole (right). Alignments of sequences were generated from mixture sequences present in the same animal and compared to the reference sequence (Ref) for the target region. The direction of sgRNA binding site shown in red arrow. CCC (surround by a red rectangle) is the PAM sequence. Three sequences are detected in the injected animal, a: wt sequence, b: sequence with nucleotides mutations around the sgRNA target sequence, c: sequence with nucleotides mutations around the sgRNA and 5 bp nucleotides deletions). **b**, Longitudinal Z-series of wholemount across the brain stem of Tg(mbp:gfp-ntr) control (upper panel) and *RetroMyelin* gRNAs-injected (RNLTR12-/-) (lower panel) Xenopus tadpoles triply labeled for GFP (green) MBP (red) and XMM (white) [XMM (*Xenopus* myelin marker): mAb recognizes specifically an unknown *Xenopus* myelin protein^65^. scale bar= 23µm. Quantification of oligodendrocytes (GFP+ cells) was plotted. P>0.05, Mann Whitney test (Two-tailed), N=4 (control), 5 (*RetroMyelin* gRNA) different sections, mean±SEM. Intensity of XMM and MBP were quantified and plotted relative to control. * P<0.05, Student’s t test (unpaired, two tailed), N=4 (control), 5 (*RetroMyelin* gRNA) different sections.

## Extended Data Table 1

*RetroMyelin* sequences in different species.

**Extended Table 1.**

| Vertebrates | Genome assembly | Top hit | E-value | Matching repeat | Family |
| --- | --- | --- | --- | --- | --- |
| Human | hg38 | chr16:35270573-35269845 | 4.6E-36 | ERV1-int (HERVS71-int) | LTR/ERV1 |
| Chimpanzee | panTro4 | chr13:18277202-18274245 | 7.4E-101 | ERV1-int (PTERV1c-int) | LTR/ERV1 |
| Gorilla | gorGor3 | CABD02423608:217-1465 | 3.6E-87 | ERV1-int (HERVS71-int) | LTR/ERV1 |
| Mouse | mm10 | chr7:30112940-30114182 | 1.5e-142 | ERV1-int (MERV1_I-int) | LTR/ERV1 |
| Rat | rn5 | chr15:24404804-24399534 | 0 | ERV1-int (RNLTR12-int) | LTR/ERV1 |
| Cow | bosTau7 | chr29:42154010-42153024 | 1.8e-29 | ERV1-int (BtERVF2_I-int) | LTR/ERV1 |
| Horse | equCab2 | chr1:27399148-27399594 | 2.9e-18 | ERV1-int (ERV1-3-EC_I-int) | LTR/ERV1 |
| Zebrafinch | taeGut1 | chr27:4207784-4209022 | 1.1e-19 | ERV1-int (TguERV2_I-int) | LTR/ERV1 |
| Chicken | galgal4 | chrW_JH375236_random:9151-8296 | 4.1E-14 | ERV1-int (GGLTR7-int) | LTR/ERV1 |
| Turtle | chrPic1 | AHGY01435164:4348-5025 | 1.3e-15 | ERV1-int (ERV1-1B_CPB-I) | LTR/ERV1 |
| Xenopus | xenLae2 | chr4S: 78506971-78506302 | 1.5e-14 | ERV1-int | LTR/ERV1 |
| Coelacanth | latCha1 | JH128184:158800-157936 | 2.5e-17 | ERV1-int | LTR/ERV1 |
| Stickleback | gasAcu1 | chrUn:56084321-56084993 | 1.7E-07 | ERV1-int<br>(Gypsy-30_GA-I) | LTR/ERV1 |
| Takifugu | fr3 | HE592075:741-39 | 4E-14 | ERV1-int<br>(FERV-R2_I-int) | LTR/ERV1 |
| Spiny chromis | ASM210954v1 | MVNR01000444.1:145194-144494 | 1.60E-08 | ERV1-int | LTR/ERV1 |
| Cichlid | NeoBri1.0 | JH422275.1:16925254-16925968 | 2.20E-10 | ERV1-int | LTR/ERV1 |
| Medaka | Om_v0.7.RACA | NVQA01006789.1:101671-102370 | 5.70E-12 | ERV1-int | LTR/ERV1 |
| Turbot | ASM318616v1 | chr4:26831079-26830372 | 4.50E-10 | ERV1-int | LTR/ERV1 |
| Atlantic cod | gadMor3.0 | chr4:29532684-29533397 | 2.1e-12 | ERV1-int | LTR/ERV1 |
| Zebrafish | danRer7 | chr15:43196012-43195219 | 1.9e-14 | ERV1-int<br>(ERV1-3-I_DR) | LTR/ERV1 |
| Whale shark | GCF_001642345.1_ASM164234v2 | NW_018046349.1:22177-21460 | 2.8e-19 | ERV1-int | LTR/ERV1 |
| Elephant shark | Callorhinchus_milii-6.1.3 | KI636671.1:5058-5765 | 1.7E-14 | ERV1-int | LTR/ERV1 |
| Lamprey | Pmarinus_7.0/petMar2 | Not found | NA | NA | NA |
| Lancelet | braflo2 | Bf_V2_65:4822224-4821722 | 3.6E-07 | (TC)n | Simple_repeat |
| Sea Urchin | strPur2 | Scaffold69942:23181-22682 | 1.70E-09 | (TC)n | Simple_repeat |
| Sea anemone | nemVec1 | Not found | NA | NA | NA |
| *C. elegans* | ce10 | Not found | NA | NA | NA |
| *Drosophila* | dm6 | Not found | NA | NA | NA |

This table list the top hit of the identified repeat type, after searching remote homology using nhmmer^41^. Specific repeat annotation, wherever available, is written under the parathesis in column 5. NA: Not applicable. Not found: wherever no hit is being obtained above the inclusion threshold (E-value<0.01).

## References

1. Zalc, B., Colman, D.R., Origins of vertebrate success. Science 288:271–2 (2000).

2. Salzer, J.L., Zalc, B., Myelination. Curr Biol. 26:R971–R975 (2016).

3. Nave, K.A., Werner, H.B., Ensheathment and Myelination of Axons: Evolution of Glial Functions. Annu Rev Neurosci. 44:197–219 (2021).

4. Lemke, G., Unwrapping the genes of myelin. Neuron 1:535–43 (1988).

5. Fields, R.D., Stevens-Graham B. New insights into neuron-glia communication. Science 298:556–62 (2002).

6. Nawaz, S., Schweitzer, J., Jahn, O., Werner, H.B. Molecular evolution of myelin basic protein, an abundant structural myelin component. Glia 61:1364–77 (2013).

7. Johnson, W.E., Origins and evolutionary consequences of ancient endogenous retroviruses. Nat Rev Microbiol 17:355–370 (2019).

8. Thompson, P.J., Macfarlan, T.S., Lorincz, M.C., Long Terminal Repeats: From Parasitic Elements to Building Blocks of the Transcriptional Regulatory Repertoire. Mol Cell 62:766–76 (2016).

9. Kazazian, H.H. Jr., Mobile elements: drivers of genome evolution. Science 303:1626–32 (2004).

10. Kazazian, H.H. Jr., Moran, J.V., Mobile DNA in Health and Disease. N Engl J Med. 377:361–370 (2017).

11. Percharde, M. et al. A LINE1-Nucleolin Partnership regulates early development and ESC identity. Cell 174:391–405 (2018).

12. Göke, J. et al. Dynamic transcription of distinct classes of endogenous retroviral elements marks specific populations of early human embryonic cells. Cell Stem Cell 16:135–41 (2015).

13. Robb, G.B., Brown, K.M., Khurana, J., Rana, T.M., Specific and potent RNAi in the nucleus of human cells. Nat Struct Mol Biol. 12:133–7 (2005).

14. Holmes, Z.E. et al. The Sox2 transcription factor binds RNA. Nat Commun 11:1805 (2020).

15. Sigova, A.A. et al. Transcription factor trapping by RNA in gene regulatory elements. Science 350:978–81 (2015).

16. Hung, T. et al., Extensive and coordinated transcription of noncoding RNAs within cell-cycle promoters. Nat Genet 43:621–9 (2011).

17. Cassiday LA, Maher LJ 3rd. Having it both ways: transcription factors that bind DNA and RNA. Nucleic Acids Res. 30:4118–26 (2002).

18. Jeon Y, Lee JT. YY1 tethers Xist RNA to the inactive X nucleation center. Cell 146:119–33 (2011).

19. Gottesfeld, J.M., Barbas, C.F. 3rd., RNA as a transcriptional activator. Chem Biol. 10:584–5 (2003).

20. Stolt, C.C. et al. Terminal differentiation of myelin-forming oligodendrocytes depends on the transcription factor Sox10. Genes Dev. 16:165–70 (2002).

21. Li, H., Lu, Y., Smith, H.K., Richardson, W.D., Olig1 and Sox10 interact synergistically to drive myelin basic protein transcription in oligodendrocytes. J Neurosci. 27:14375–82 (2007).

22. Bourque, G. et al. Ten things you should know about transposable elements. Genome Biol. 19:199 (2018).

23. Goldman, N., Whelan, S., Statistical tests of gamma-distributed rate heterogeneity in models of sequence evolution in phylogenetics. Mol Biol Evol. 17:975–8 (2000).

24. McKenzie, I.A., et. al. Motor skill learning requires active central myelination. Science 346:318–22 (2014).

25. Xin, W., Chan, J.R., Myelin plasticity: sculpting circuits in learning and memory. Nat Rev Neurosci. 21:682–694 (2020).

26. Franklin, R.J.M., ffrench-Constant, C., Regenerating CNS myelin - from mechanisms to experimental medicines. Nat Rev Neurosci. 18:753–769 (2017).

27. Fancy, S.P., Chan, J.R., Baranzini, S.E., Franklin, R.J.M., Rowitch, D.H., Myelin regeneration: a recapitulation of development? Annu Rev Neurosci. 34:21–43 (2011).

## References

28. Reichmann, J., et al. Microarray analysis of LTR retrotransposon silencing identifies Hdac1 as a regulator of retrotransposon expression in mouse embryonic stem cells. PLoS Comput Biol. 8:e1002486 (2012).

29. AFA Smit, R Hubley and P Green, RepeatMasker Open-4.0. 2013-2015 http://www.repeatmasker.org.

30. Ritchie, M.E. et al. limma powers differential expression analyses for RNA-sequencing and microarray studies. Nucleic Acids Res. 43:e47 (2015).

31. Langfelder, P., Horvath, S., WGCNA: an R package for weighted correlation network analysis. BMC Bioinformatics 9:559 (2008).

32. Becker RA, et al., The New S Language. Wadsworth & Brooks/Cole (1988).

33. Huang, da W., Sherman, B.T., Lempicki, R.A., Systematic and integrative analysis of large gene lists using DAVID bioinformatics resources. Nat Protoc. 4:44–57 (2009).

34. Guo, J.C. et al. CNIT: a fast and accurate web tool for identifying protein-coding and long non-coding transcripts based on intrinsic sequence composition. Nucleic Acids Res. 47:W516–W522 (2019).

35. Kopylova E, Noé L, Touzet H. SortMeRNA: fast and accurate filtering of ribosomal RNAs in metatranscriptomic data. Bioinformatics 28:3211–7 (2012).

36. Bolger, A.M., Lohse, M., Usadel, B., Trimmomatic: a flexible trimmer for Illumina sequence data. Bioinformatics 30:2114–20 (2014).

37. Dobin, A. et al. STAR: ultrafast universal RNA-seq aligner. Bioinformatics 29:15–21 (2013).

38. Li, H. et al. 1000 Genome Project Data Processing Subgroup. The sequence alignment/map format and SAMtools. Bioinformatics 25:2078–9 (2009).

39. Patro, R., Duggal, G., Love, M.I., Irizarry, R.A., Kingsford, C., Salmon provides fast and bias-aware quantification of transcript expression. Nat Methods 14:417–419 (2017).

40. Love, M.I., Huber, W., Anders, S., Moderated estimation of fold change and dispersion for RNA-seq data with DESeq2. Genome Biol. 15(12):550 (2014).

41. Wheeler, T.J., Eddy, S.R., nhmmer: DNA homology search with profile HMMs. Bioinformatics 29:2487–9 (2013).

42. Letunic, I., Bork, P., Interactive Tree Of Life (iTOL) v4: recent updates and new developments. Nucleic Acids Res. 47:W256–W259 (2019).

43. Katoh, K., Standley, D.M., MAFFT multiple sequence alignment software version 7: improvements in performance and usability. Mol Biol Evol. 30:772–80 (2013).

44. Criscuolo, A., Gribaldo, S., BMGE (Block Mapping and Gathering with Entropy): a new software for selection of phylogenetic informative regions from multiple sequence alignments. BMC Evol Biol. 10:210 (2010).

45. Stamatakis, A., RAxML version 8: a tool for phylogenetic analysis and post-analysis of large phylogenies. Bioinformatics 30:1312–3 (2014).

46. Pattengale, N.D. et al. How many bootstrap replicates are necessary? J Comput Biol. 17:337–54 (2010).

47. Yang, Z., PAML: a program package for phylogenetic analysis by maximum likelihood. Comput Appl Biosci. 13:555–6 (1997).

48. Liberles, D.A., Dittmar, K., Characterizing gene family evolution. Biol Proced Online 10:66–73 (2008).

49. Yang, Z., Estimating the pattern of nucleotide substitution. J Mol Evol. 39:105–11 (1994).

50. Yang, Z., Maximum likelihood phylogenetic estimation from DNA sequences with variable rates over sites: approximate methods. J Mol Evol. 39:306–14 (1994).

51. Yang, Z., Wang, T., Mixed model analysis of DNA sequence evolution. Biometrics 51:552–61 (1995).

52. Segel, M. et al. Niche stiffness underlies the ageing of central nervous system progenitor cells. Nature 573:130–134 (2019).

53. Neumann, B. et al. Metformin restores CNS remyelination capacity by rejuvenating aged stem cells. Cell Stem Cell 25:473–485 (2019).

54. Pol, S.U. et al. Sox10-MCS5 enhancer dynamically tracks human oligodendrocyte progenitor fate. Exp Neurol 247:694–702 (2013).

55. Rawji, K.S. et al. Deficient surveillance and phagocytic activity of myeloid cells within demyelinated lesions in aging mice visualized by ex vivo live multiphoton imaging. J Neurosci. 38:1973–1988 (2018).

56. Schindelin, J., et al. Fiji: an open-source platform for biological-image analysis. Nat Methods 9:676–82 (2012).

57. Pfaffl, M.W., A new mathematical model for relative quantification in real-time RT-PCR. Nucleic Acids Res. 29:e45 (2001).

58. Weinmann, A.S., Farnham, P.J., Identification of unknown target genes of human transcription factors using chromatin immunoprecipitation. Methods 26:37–47 (2002).

59. Ghosh, T. et al. MicroRNAs establish robustness and adaptability of a critical gene network to regulate progenitor fate decisions during cortical neurogenesis. Cell Rep. 7:1779–88 (2014).

60. Almeida, R.G., Czopka, T., Ffrench-Constant, C., Lyons, D.A., Individual axons regulate the myelinating potential of single oligodendrocytes in vivo. Development 138:4443–50 (2011).

61. Early, J.J. et al. An automated high-resolution in vivo screen in zebrafish to identify chemical regulators of myelination. Elife 7:e35136 (2018).

62. Meeker, N.D., Hutchinson, S.A., Ho, L., Trede, N.S., Method for isolation of PCR-ready genomic DNA from zebrafish tissues. Biotechniques 43:610–614 (2007).

63. Mannioui, A. et al. The Xenopus tadpole: An in vivo model to screen drugs favoring remyelination. Mult Scler. 24:1421–1432 (2018).

64. Dehairs, J., Talebi, A., Cherifi, Y., Swinnen, J.V., CRISP-ID: decoding CRISPR mediated indels by Sanger sequencing. Sci Rep. 6:28973 (2016).

65. Steen, P., Kalghatgi, L., Constantine-Paton, M., Monoclonal antibody markers for amphibian oligodendrocytes and neurons. J Comp Neurol. 289:467–80 (1989).

66. Royston, J.P., Appl. Stat. 181:176–180 (1982).

